# Hierarchical communities in the larval *Drosophila* connectome: Links to cellular annotations and network topology

**DOI:** 10.1101/2023.10.25.562730

**Authors:** Richard Betzel, Maria Grazia Puxeddu, Caio Seguin

## Abstract

One of the longstanding aims of network neuroscience is to link a connectome’s topological properties–i.e. features defined from connectivity alone–with an organism’s neurobiology. One approach for doing so is to compare connectome properties with maps of metabolic, functional, and neurochemical annotations. This type of analysis is popular at the meso-/macro-scale, but is less common at the nano-scale, owing to a paucity of neuron-level connectome data. However, recent methodological advances have made possible the reconstruction of whole-brain connectomes at single-neuron resolution for a select set of organisms. These include the fruit fly, *Drosophila melanogaster*, and its developing larvae. In addition to fine-scale descriptions of neuron-to-neuron connectivity, these datasets are accompanied by rich annotations, documenting cell type and function. Here, we use a hierarchical and weighted variant of the stochastic blockmodel to detect multi-level communities in a recently published larval *Drosophila* connectome. We find that these communities partition neurons based on function and cell type. We find that communities mostly interact assortatively, reflecting the principle of functional segregation. However, a small number of communities interact non-assortatively. The neurons that make up these communities also form a “rich-club”, composed mostly of interneurons that receive sensory/ascending inputs and deliver outputs along descending pathways. Next, we investigate the role of community structure in shaping neuron-to-neuron communication patterns. We find that polysynaptic signaling follows specific trajectories across modular hierarchies, with interneurons playing a key role in mediating communication routes between modules and hierarchical scales. Our work suggests a relationship between the system-level architecture of an organism’s complete neuronal wiring network and the precise biological function and classification of its individual neurons. We envision our study as an important step towards bridging the gap between complex systems and neurobiological lines of investigation in brain sciences.

## INTRODUCTION

Nervous systems are fundamentally networks [1, 2]. They are composed of neural elements–cells, areas, regions–linked to one another *via* synapses, axonal projections, and myelinated white matter. The complete set of neural elements and their pairwise connections defines a “connectome”. The configuration of the connectome as a network helps shape brain activity and function.

A popular strategy for analyzing connectome data is to represent it as a graph of nodes and edges [3]. This simple model generally abstracts away neurobiological detail, but returns the backbone of structural interactions, which can be further analyzed using network science tools. Network science sits at the confluence of statistical physics, engineering, and mathematics, and offers a wide range of tools for summarizing and characterizing the structure and function of connectome data.

In principle, the network model is agnostic to scale; it is equally well-suited for representing large-scale connectivity (regions/areas linked by fiber tracts/projections) [4–6] as it is for representing cellular-level connectivity (synaptically coupled neurons) [7]. Network analyses have identified a number of phylogenetically conserved architectural features of connectomes, including efficient processing paths coupled with greater-than-expected clustering (small-worldness [8–11 heterogeneous degree distributions and inter-linked hubs [4, 12, 13], cost-efficient spatial embedding [14– and neurobiologically meaningful sub-networks or modules [17, 18].

To date, however, most connectome analyses have focused on the macroscale, as data can be acquired cheaply, non-invasively (diffusion magnetic resonance imaging + tractography), and for the entire brain at a single-subject level. Indeed, very few connectome datasets exist at both the whole-brain and single-cell levels – the most notable being that of the nematode *C. elegans* [7, 19, 20]. Recently, however, methodological advances have made it possible to reconstruct cellular-level connectivity for large volumes [21–28]. Importantly, reconstructions of synaptic connectivity are accompanied by rich, high-dimensional sets of neurobiological annotations, which are seldom available for macroscale connectomes. These include details of cells’ morphologies, types and lineages, as well as putative functional assignments, thus making it possible to link high-level architectural features to fine-grained properties of single neurons at the same spatial resolution where neurobiological processes unfold.

Here, we apply network science tools to the connectome of the larval *Drosophila melanogaster*. Our focus is on its community structure, which we uncover using an extension of the classical stochastic blockmodel [29, 30]. We find evidence of hierarchical community structure – communities within communities – whose sizes range from tens to hundreds of neurons. Community boundaries sharply delineate different cell types from one another and circumscribe groups of cells with shared functional profiles, e.g. proprioception, nociception, olfaction, and so on. Further, we show that while most communities interact assortatively–i.e. form internally dense and externally sparse subnetworks–a small fraction form nonassortative motifs and that these communities are enriched for interneurons. Next, we show that the larval *Drosophila* brain exhibits a “rich club”– a collection of hubs that are also mutually connected to one another. We show that rich-club neurons, which tend to be interneurons, are present in most coarse-scale modules, but are also concentrated within a select set of “hub” modules, all positioned in the midbrain. Lastly, we investigated the link between community structure and communication policies, focusing on shortest paths and communicability. In general, we found that neurons that were frequently assigned to the same community across hierarchical levels were also more likely to have greater pairwise communicability and reduced path length. Collectively, these results recapitulate, in a whole-brain nano-scale connectome, a set of architectural features that are largely conserved across phylogeny and scale.

## RESULTS

Here, we analyze a previously published nano-scale connectome for the *Drosophila melanogaster*. The connectome is comprised of *N* = 2952 cells and approximately *K* = 352611 synapses.

### Hierarchical community structure

Connectomes are thought to be modular, meaning that they can be decomposed into meaningful subgraphs referred to as “communities”. Here, we use a hierarchical variant of the stochastic blockmodel (SBM) to partition the connectome into nested communities [30]. Unlike modularity maximization [31] or Infomap [32], which are the most popular community detection methods in network neuroscience but only capable of detecting assortative community structure (internally dense and externally sparse subgraphs), SBMs use an inferential framework to detect generalized classes of communities, including core-periphery and disassortative motifs [29, 30, 33–35] (though note that the SBM can be constrained to detect purely assortative partitions [36]). In addition, the SBM does not suffer from overfitting issues that permit methods like modularity maximization and Infomap to detect communities in random networks [37].

To obtain an estimate of hierarchical communities, we used the procedure described in Peixoto [38] to sample a large number of high-quality partitions (10000 samples) and discover latent modes. We found evidence supporting the hypothesis that there is exactly one partition mode. Here, we characterize the consensus estimate of that partition. The optimal partition resulted in seven hierarchical levels, dividing the network into 2, 4, 6, 10, 20, 36, and 77 communities (Fig. 1*a-c*). For the sake of visualization, we focus on the fourth hierarchical level. In Fig. 1*d* we visualize these communities, coloring neurons and their morphological trees based on the community to which they were assigned. We note, however, that these ten communities can be both sub-divided and aggregated further. We show in Fig. 1*e* examples of communities in the fourth hierarchical level that fracture into two or three smaller communities in the fifth level. For alternative visualizations of the different hierarchical levels and for a qualitative assessment of the link to synapse type-specific connectomes, see Fig. S1 and Fig. S2, respectively.

**Figure 1.**
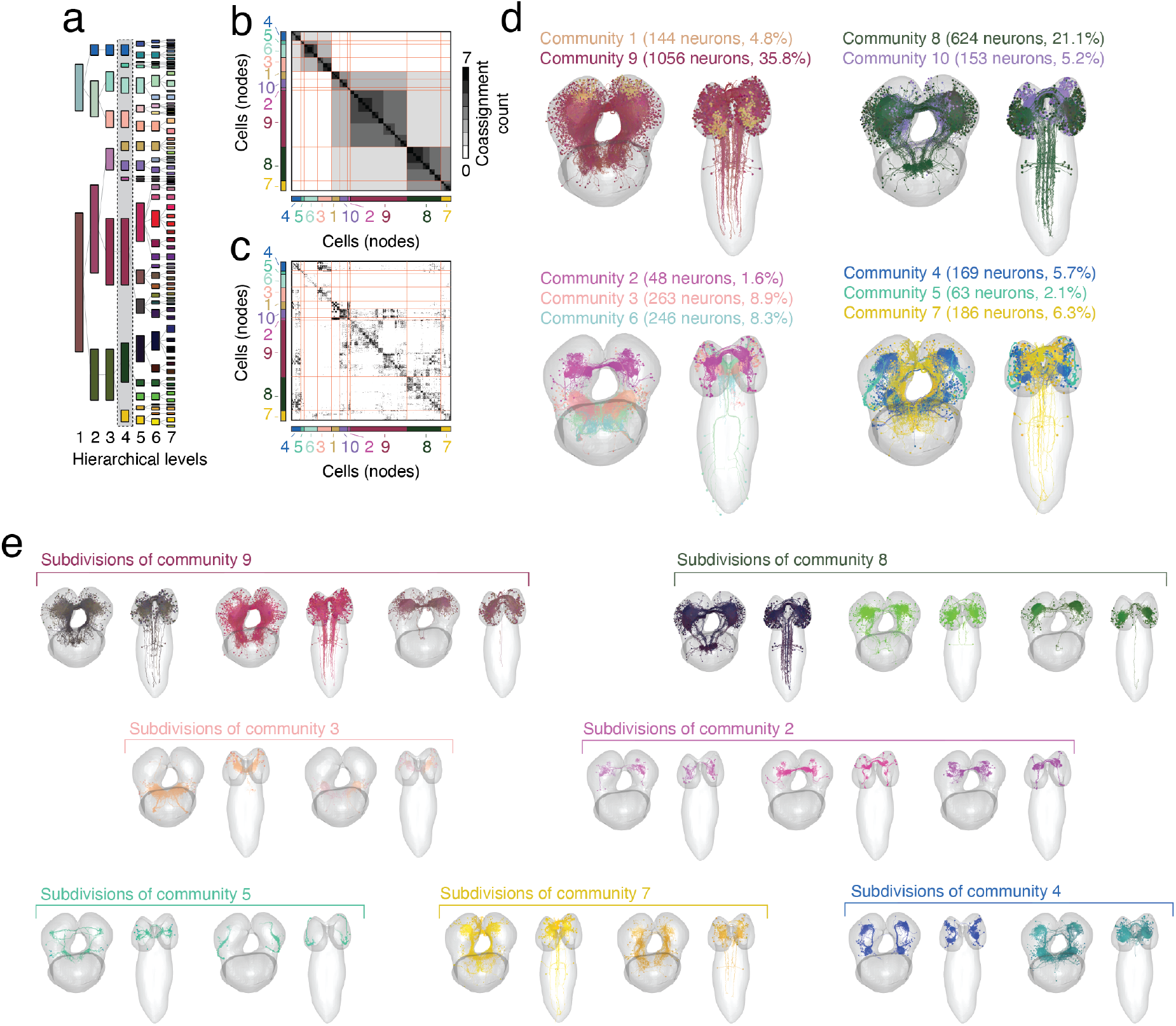
Detected hierarchical modular structure. (*a*) Hierarchical dendrogram. Each column corresponds to a hierarchical level. Colors correspond to communities at the finest hierarchical level (level five). At coarser levels they are grouped into larger communities. (*b*) Community co-assignment matrix. Entries correspond to the number of levels in which pairs of neurons were assigned to the same community. Red lines separate level four communities from one another. (*c*) Connectivity matrix with rows/columns ordered by communities. Panel *d* depicts level-four communities in anatomical space. Communities can, in general, be subdivided or even aggregated into larger clusters. Panel *e* highlights divisions of select communities in the fourth hierarchical level into smaller sub-communities in level five.

Next, we aimed to characterize the profile of communities across hierarchical levels. As expected, the size of communities–the number of nodes assigned to a given community–decreased monontonically with hierarchical level (Fig. 2*a*). In parallel, the synaptic and binary densities of communities increased across hierarchical levels (Fig. 2*b,c*), suggesting that these communities become more internally cohesive.

**Figure 2.**
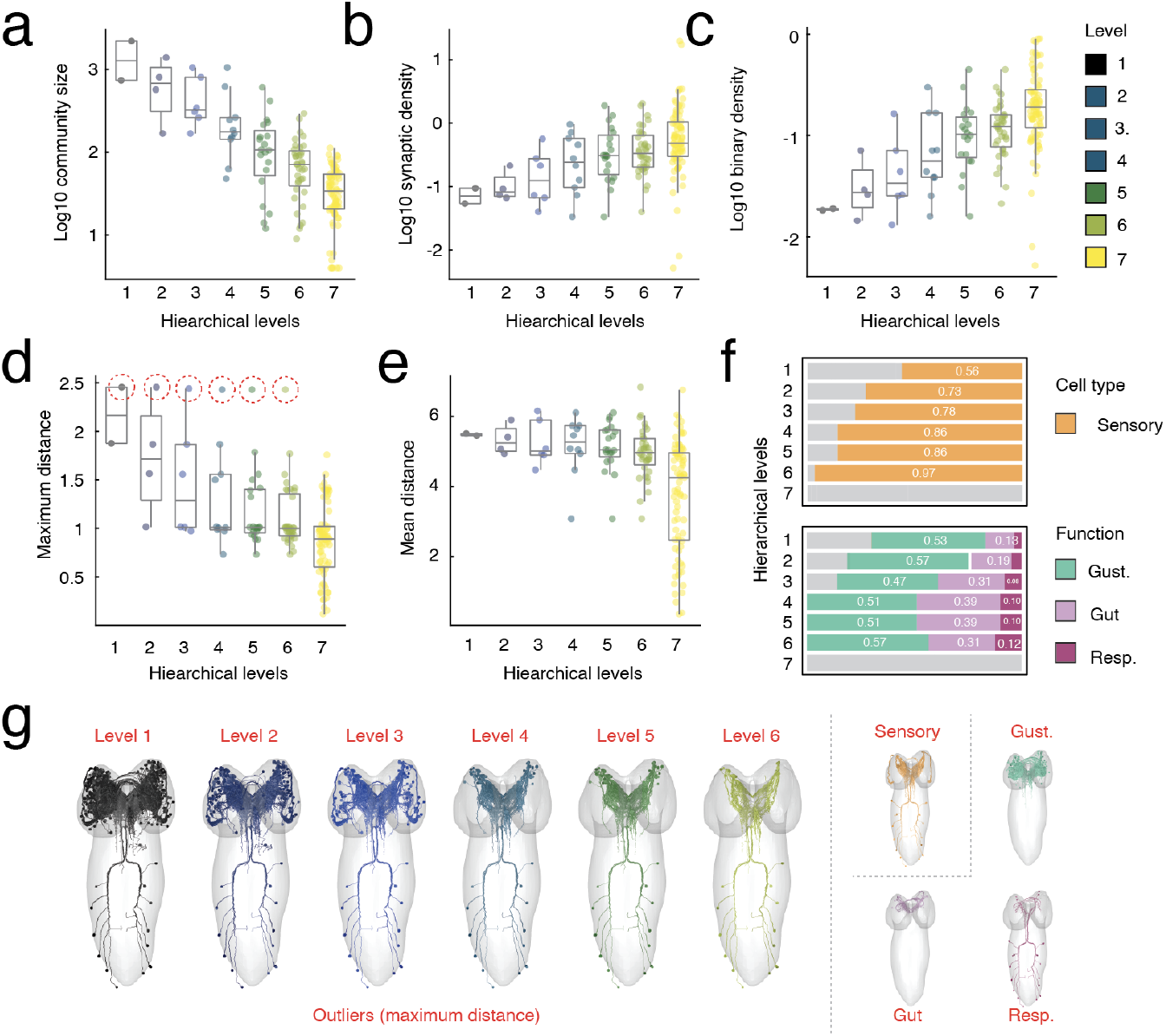
Community statistics across hierarchical levels. (*a*) Community size (number of nodes). (*b*) Log synaptic density – number of synapses divided by total number of possible connections within each community. (*c*) Log binary density – number of connections divided by total number of possible connections. (*d*) Community diameter – longest distance (Euclidean) between pairs of nodes assigned to the same community. (*e*) Mean distance between pairs of nodes. At each hierarchical level there were clear outliers in terms of maximum distance (red circles in panel *d*). In panel *f*, we break down the cell type and functional composition of those communities. In panel *g* we show the anatomical configuration of those communities. In general, they are composed of ascending sensory projections that support gustatory, gut, and respiratory function.

Next, we examined the spatial properties of communities. We found that the mean and maximum pair-wise (Euclidean) distance between soma decreased near-monotonically across hierarchical levels (Fig. 2*d,e*). We note, however, that there were several outliers – communities whose diameter was far greater than the typical community. These communities were composed largely of sensory neurons in the nerve cord associated with respiratory, gut, and gustatory function (Fig. 2*f,g*).

In the **Supplementary Material** we investigated other generic spatial properties of the larval *Drosophila* connectome. One of the organizing principles of inter-areal connectomes is the exponential distance rule (EDR), wherein connection weight decays approximately as an exponential function with inter-areal Euclidean distance [39, 40]. However, whether this rule holds at the nano-scale remains largely untested. To address this question, we examined how the distances between cells were related to their synapse counts (see Fig. S3). When the EDR is described in inter-areal connectome studies, distance is usually operationalized as the Euclidean distance between areal centers of mass. Accordingly, we considered soma-soma distance as one distance metric. However, synaptic contacts between neurons in the larval *Drosophila* are generally distant from their respective soma, such that two spatially adjacency cells (if they connect) do so a long distance away from their cell bodies. Accordingly, we also defined a second distance metric that takes into account the location of synapses. Specifically, for a pre- and post-synaptic neuron, we calculated the shortest path through their arbors from their respective soma to location of the synapse. We then calculated the wiring cost along the shortest path as a measure of distance. If two neurons were connected *via* multiple synapses, we identified the union of their shortest paths (the shortest paths backbone) and calculated the Euclidean distance along all arboral segments comprising said backbone.

See Fig. S4 for a direct comparison of these two measures.

In general, we found that although connection probability decayed with soma-soma distance (Fig. S3e), synapse count was not correlated with soma-soma Euclidean distance (*r* = −0.0015; Fig. S3c) but was positively correlated with the shortest path distance (*r* = 0.24; *p <* 10^−15^; Fig. S3d). Interestingly, we found that the strength of association between synapse count and the wiring cost along the shortest path was cell-type specific (Fig. S3g-i).

These results suggest that a simple exponential distance rule may not hold at the level of synaptic connectivity. However, if we coarse-grain the network by partitioning neurons into clusters, we find that connection weights in the cluster-level representations of the network exhibit negative relationship with distance – both soma-soma distance and wiring cost along the shortest path – that is modulated by the level of granularity. For very coarse representations–tens or hundreds of clusters–we observe strong anticorrelations (Fig. S3f). However, as *k* → *N*, the correlation magnitude increases, approaching 0 in the case of soma-soma distance and ≈ 0.24 in the case of distance along the shortest path. These observations suggest that, at single-neuron resolution, the effect of straight-line distance on synapse count is small and possibly inappropriate – i.e. there are better measures of distance that *are* predictive of synapse count [41].

In addition to the SBM, we also tested a hierarchical variant of modularity maximization, the results of which we report in the supplementary material (Fig. S5). Note further that, despite differences in objective function and optimization heuristic, the communities detected using the SBM and modularity maximization are highly similar (Fig. S6), yielding correlated enrichment scores (Fig. S7). We also compared the partitions obtained using the SBM with a subset of the partitions reported in Winding *et al*. [21] (Fig. S10). In general, we found that our solutions were neither a perfect alignment with the previously reported clusters, nor were they wildly dissimilar. Lastly, we also fit the connectomes comprising only axon→axon, axons→dendrite, dendrite→axon, and dendrite→dendrite connectomes (Fig. S11). Overall, we found that communities inferred from the axon→axon and axons→dendrite connectomes were most similar to the results described here and that the dendrite→dendrite connectome exhibited marked community laterality–i.e. communities tended to be comprised of neurons in one or the other hemisphere.

### Linking communities to cell types and function

In the previous section, we described a set of hierarchical modules and their properties. How do these modules and their boundaries relate to cellular annotations? Do they “carve nature at its joints” such that modules circumscribe specific types of cells or functions? Or are annotations intermixed evenly across modules? At the meso-/macro-scale this question can only be approximately addressed by averaging cellular or population-level annotations to generate parcel-based maps [42]. Here, however, we take advantage of the fact that communities and annotations are both defined at the single-neuron scale, allowing us to compare them directly. Specifically, we asked whether connectivity-defined communities were preferentially “enriched” for different types of annotations: neurons associations with specific functions and cell types, as well as broader classes of neurons (e.g. input, output, interneuron) [43]. We calculated the overlap of each annotation and community–e.g. the number of nodes labeled as Kenyon cells that were also assigned to community *C*. We then compared the observed overlap against a null distribution generated under a null model that preserves the spatial variogram [44] (Fig. 3*a*).

**Figure 3.**
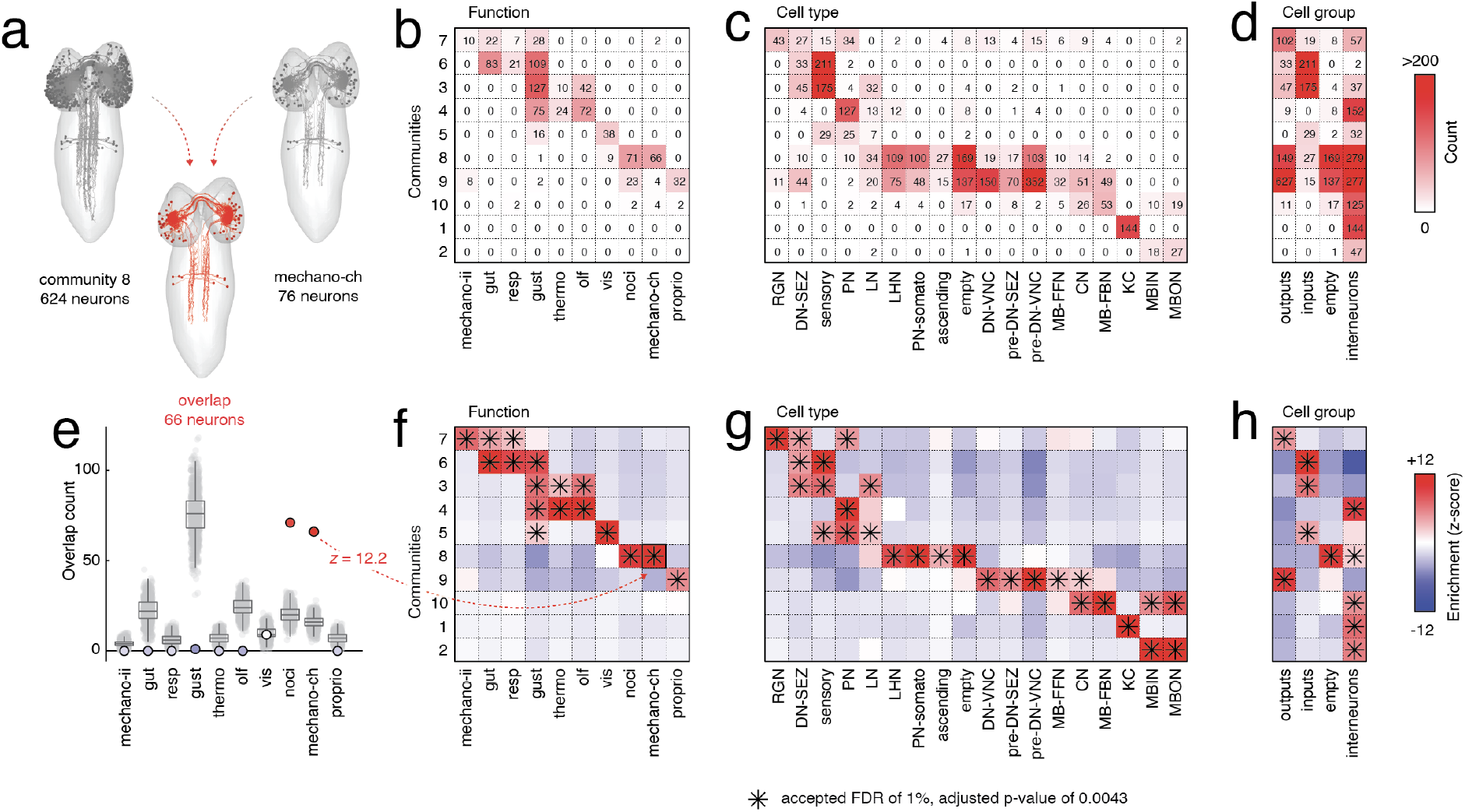
Communities are “enriched” for function, cell type, and cell group. (*a*) Schematic illustrating how we measure overlap. Given two partitions of cells – e.g. one coming from data-driven community labels and another coming from annotation data – we calculate the overlap as the union of the two. Panels *b*-*d* show the overlap (counts) for each community and for different functional groups, cell types, and macro cell group labels. (*e*) The counts are, in general, confounded by community and annotation size – larger maps will, just by chance, tend to have greater overlap with one another. To control for this, we use a space-preserving null model to calculate the expected overlap for each entry in the arrays depicted in panels *b*-*d*, and z-score the observed overlap scores with respect to the null distributions. Panel *e* illustrates this procedure. This allows us to contextualize the scores shown in panels *b*-*d* – entries with greater overlap than expected signify that a community may be “enriched” for a given function, cell type, or macro cell group.

We found that communities exhibited both a high level of enrichment and also a high degree of specificity in terms of the annotations for which they were enriched (see Fig. 3*b* for raw overlap scores and Fig. 3*f* for raw scores normalized against null the null distribution and expressed as z-scores). This is interesting, as communities were defined only on the basis of synaptic connectivity using a data-driven algorithm. Virtually every function had a clear correspondence with a community. Community 9 was associated with proprioception; community 8 was dually associated with nociception and chordotonal mechanosensation; community 5 was associated with vision and gustation; communities 3 and 4 were jointly associated with gustation, thermosensation, and olfaction; community 6 wassociated with gut, respiration, and gustation; community 7 was associated with class II mechanosensation, gut function, and respiration. Interestingly, communities 1, 2, and 10, which were among the smallest and spatially compact, were composed mostly of interneurons and not clearly aligned with any sensorimotor function.

Similarly, cell types (and by extension, the broader cell classes/groups to which cell types were assigned) were also significantly enriched within communities. For example, communities 1, 2, and 10, which were poorly aligned with functional annotations, were highly enriched for different classes of interneurons, with community 1 aligned with Kenyon cells, community 2 aligned with mushroom body input/output neurons, and community 10 aligned with mushroom body input/output/feedback neurons (see Fig. 3*c,d* for raw overlap; Fig. 3*g,h* for z-scored enrichment scores).

We also linked connectional properties of communities with function, class, and cell type. Specifically, we calculated “community motifs”, which describe how pairs of communities interact with one another [34]. Every pair of communities can be represented as a [2 *×* 2] matrix, whose diagonal elements correspond to the within-community synaptic densities and the off-diagonal elements correspond to the synaptic density of incoming/outgoing connections. Based on these four elements, the interactions between every pair of communities can be unambiguously classified as either assortative, core-periphery, or disassortative (Fig. 4*a,b*).

**Figure 4.**
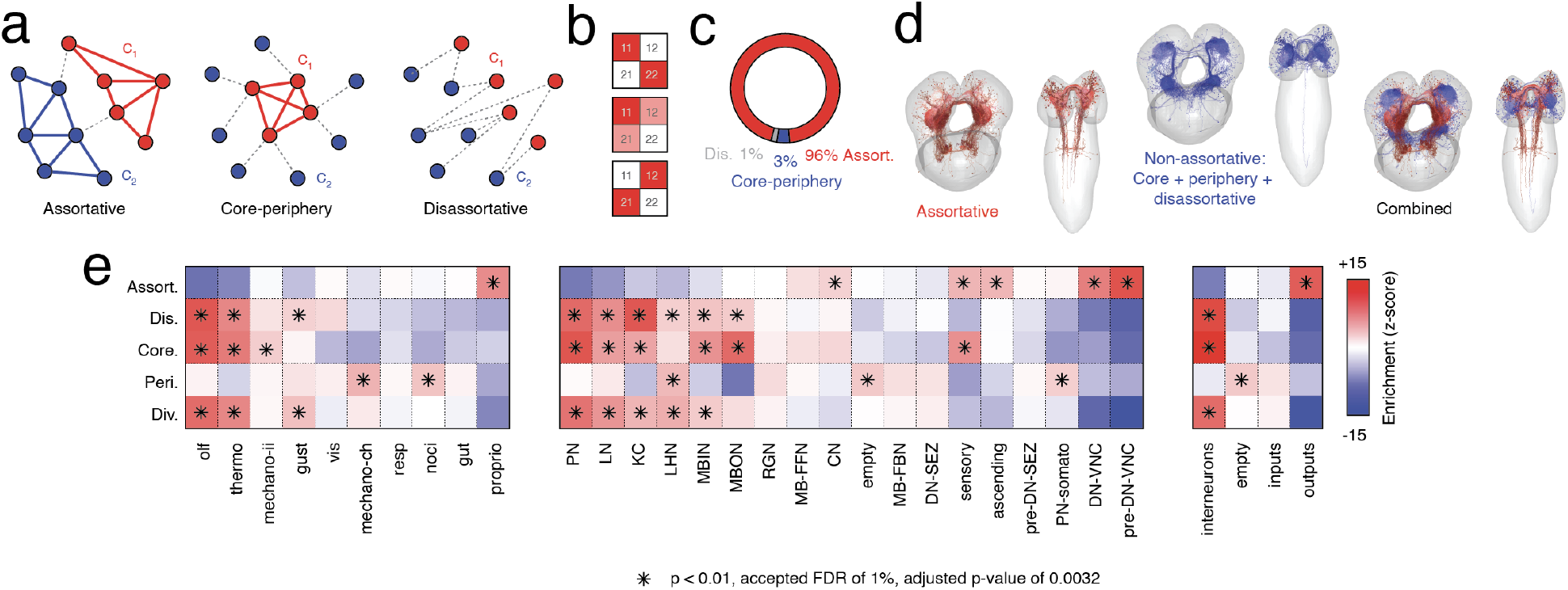
Community motifs distinguish function, cell type, and macro cell group from one another. (*a*) Example community interaction motifs. Assortative interactions correspond to “internally dense - externally sparse” communities; “core-periphery” interactions involve a densely connected core that is linked to a sparse periphery; “disassortative” communities form cross-community links, seldom connecting nodes of the same community to one another. (*b*) Density matrices for the community interaction motifs discussed in *a*. (*c*) Proportion of all two-community motifs classified as either assortative, core-periphery, or disassortative. (*d*) Anatomical depiction of the most assortative and non-assortative neurons and their arbors. (*e*) Enrichment (z-score) of assortative, disassortative, core, periphery, and diversity indices within function, cell type, and macro cell group labels. Panels *f, g*, and *h* report “enrichment scores” (z-scores) of the raw counts depicted in panels *b, c*, and *c*.

Here, we categorize the interaction between every pair of communities and map these labels back to individual network nodes. This procedure is carried out independently for each hierarchical level, the scores ranked, and then averaged across levels to yield a single assortativity, core, periphery, and disassortativity score per neuron. Note that for core-periphery motifs, we distinguish between the community that acts as the “core” and the community that acts as the “periphery”. Based on these labels, we also calculated an entropy-based “diversity index” whose value is close to 1 if a node participates uniformly in all four classes and is close to 0 if it participates in only a single class. For the sake of visualization, we group the non-assortative labels (core, periphery, and disassortative) together to form a “non-assortative” class (Fig. 4*c*). We then performed another enrichment analysis, this time testing whether community motif values were significantly concentrated within functional, cell type, and cell class groups.

We found, in line with other applications of the stochastic blockmodel to brain network data, that most communities interact assortatively [34, 35]. Specifically, 96% of interactions are assortative, with only 3% and 1% classified as core-periphery and disassortative, respectively (Fig. 4*c,d*). Interestingly, we found that motif classes were enriched within functional groups, cell types, and cell classes. For example, assortative motifs were highly enriched within output neurons, namely descending neurons in the ventral nerve cord associated with proprioception, nociception, and chordotonal mechanosensation (Fig. 4*e*). Note that this analysis only implies that the level of assortativity within these groups was greater than expected by chance; it does not imply that these groups were the only groups that participated in assortative community motifs. Conversely we found that interneurons specifically were enriched for the non-assortative community motifs–i.e. disassortativity and coreness–and were, in general, among the most diverse in terms of their motif-type participation (Fig. 4*d,e*).

Collectively, these results support the hypothesis that the *Drosophila* larval connectome exhibits hierarchical modular structure. These communities tend to be assortative and, though data-driven, divide cells into groups based on their type, class, and function. Further, communities occasionally deviate from assortative interactions; the cells that make up these communities tend to be interneurons, supporting the hypothesis that the most assortative and segregated communities support specialized brain function.

### Rich-club structure

Another hallmark feature of brain networks is that their degree distributions tend to be heavy-tailed, signifying that most neural elements maintain few partners, but that a small number make disproportionately many connections. In other brain networks datasets, these highly connected nodes sometimes form a so-called “rich-club” [45– wherein hub nodes are more densely connected to one another than expected by chance [12, 13]. Rich-clubs are thought to be essential features for inter-modular communication–human studies have found that rich-club nodes, though relatively rare, are distributed across cortical modules [48], though other studies using blockmodels to define communities have shown that rich-clubs can form their own, separate community [49]. Here, we test whether the larval *Drosophila* connectome exhibits a rich-club and, if so, assess how it inter-links communities to one another, as well as its relationship to known functional classes and cell types.

Specifically, we calculated the directed rich-club coefficient, *ϕ*(*k*), across all possible values of total degree, *k*. We then repeated this procedure for 1000 randomized networks whose incoming and outgoing degree sequences were identical to that of the original network but where the connections were otherwise formed at random [50]. Then, for each value of *k*, we calculated the non-parametric *p*-value as the fraction of randomized networks whose rich-club coefficient was equal or greater to that of the original network (Fig. 5*a*). We identified a range of statistically significant rich-clubs, but focused on the local maxima in the normalized rich-club coefficient at *k* = 172.

**Figure 5.**
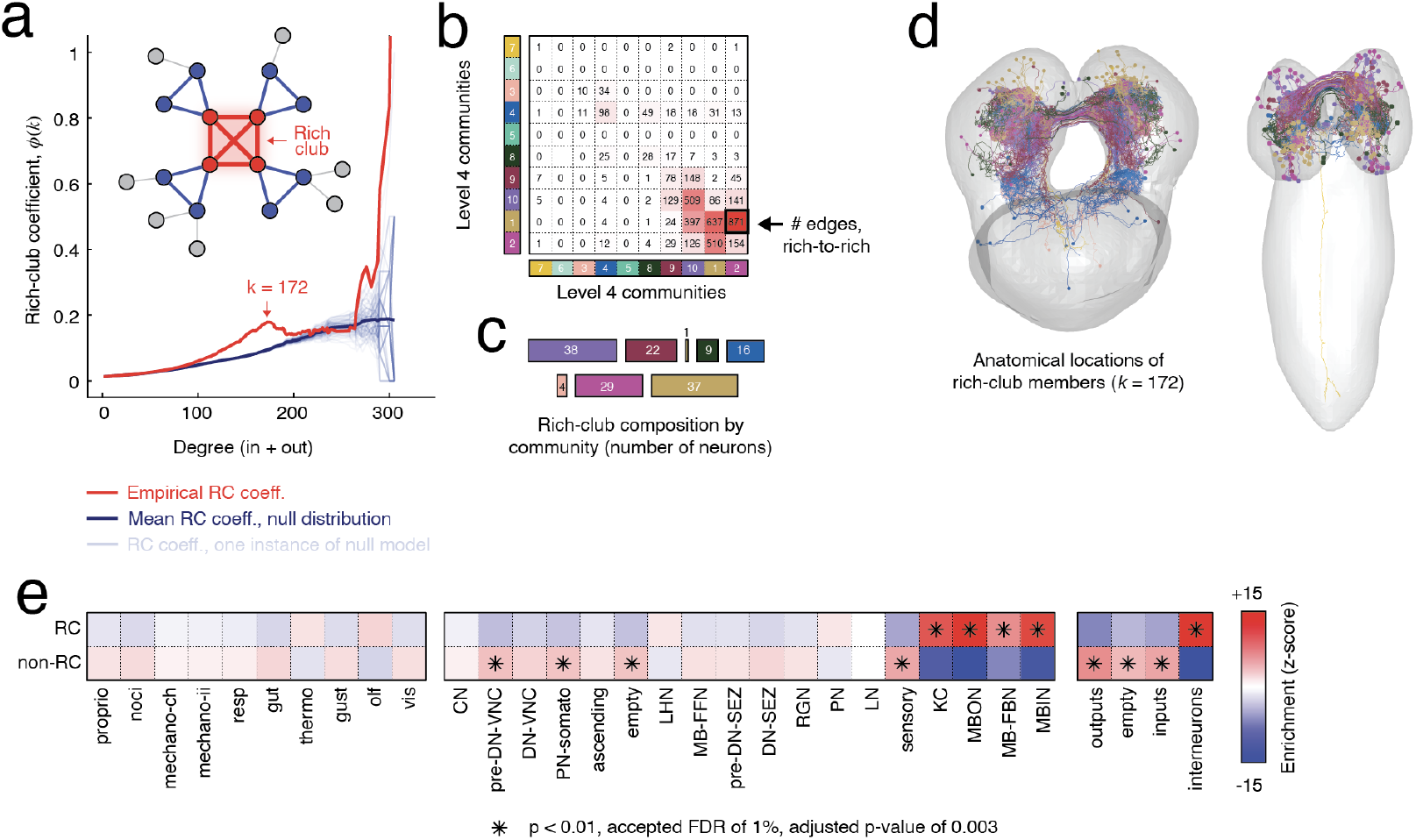
Rich-club structure. (*a*) Empirical rich-club coefficient (red) compared to randomized null models (blue). We focus on the statistically significant rich-club at *k* = 172 (combined in-/out-degree). (*b*) Concentration of rich-club connections (connections that link a rich-club node to another rich-club member. (*c*) Composition of rich-club by communities. Colors correspond to level two communities; area is proportional to the percentage of the rich-club that comes from each community. (*d*) Anatomical representation of rich-club nodes and their arbors. (*e*) Enrichment of function, cell type, and macro cell group within rich-club labels (part of rich-club and not part of rich-club).

We found that rich-club nodes were distributed across 8/10 communities at the fourth hierarchical level (albeit only barely; community 7 included only 1 rich-club node rich-club connection) (Fig. 5*b-d*). However, rich-club nodes were largely concentrated within four communities (1, 2, 9, and 10), whose constituent members were among the most connected in the network as indexed by degree (Fig. S8). We also tested whether rich-club membership was linked to specific function, cell types, or cell groups using the same enrichment analysis described in the previous section. We found that interneurons, specifically Kenyon cells and a subset of cells in the mushroom body (input, output, and feedback neurons [51]), were preferentially associated with rich-club status (*p <* 10^−15^; Fig. 5*e*).

We also used information about synapse type to further decompose and characterize the rich-club (Fig. S12). In general, we found that “rich” connections (links between two rich-club neurons) were most likely to reflect axon→axon and axon→dendrite synapses, reflecting the fact that these two synapse types were also the most common (Fig. S12b). However, when we controlled for baseline rate, we found that axon→axon synapses were significantly overrepresented while axon→dendrite synapses were significantly under-represented (permutation test; *p <* 0.01). Interestingly, for more exclusive rich-clubs (*k >* 172) we find evidence that dendrite→axon and axon→dendrite synapses are slightly overrepresented relative to their baseline rate (Fig. S12c), suggesting that these synapses may be important for signaling amongst highly connected, hub neurons.

Collectively, these results indicate that the *Drosophila* connectome exhibits rich-clubs – groups of highly-connected cells that are also connected to one another. Here, the rich-club was detected in a data-driven way but overlaps with known cell types that have been linked to associative learning and memory [52].

### Role of network modules in communication processes

Connectomes represent the pathways along which signals propagate. Communication between cells can be understood as the process by which a “message” or “signal” from a source node reaches a pre-specified target [53, 54]. These types of processes can be modeled using tools from network science. Typically, communication models are situated along a spectrum, ranging from centralized processes like shortest paths routing, in which the message follows the shortest possible path from its source to target, ensuring maximum efficiency, to decentralized processes like diffusion [55, 56], navigation [57], and cascade models [58]. Here, we explore communication processes unfolding over the *Drosophila* connectome. Although the set of possible communication policies is vast [59, 60], we restrict our analyses to the following three:

#### 1. Binary shortest paths

the fewest number of hops between pairs of neurons if we were to discard edge weights (Fig. 6*a*),

**Figure 6.**
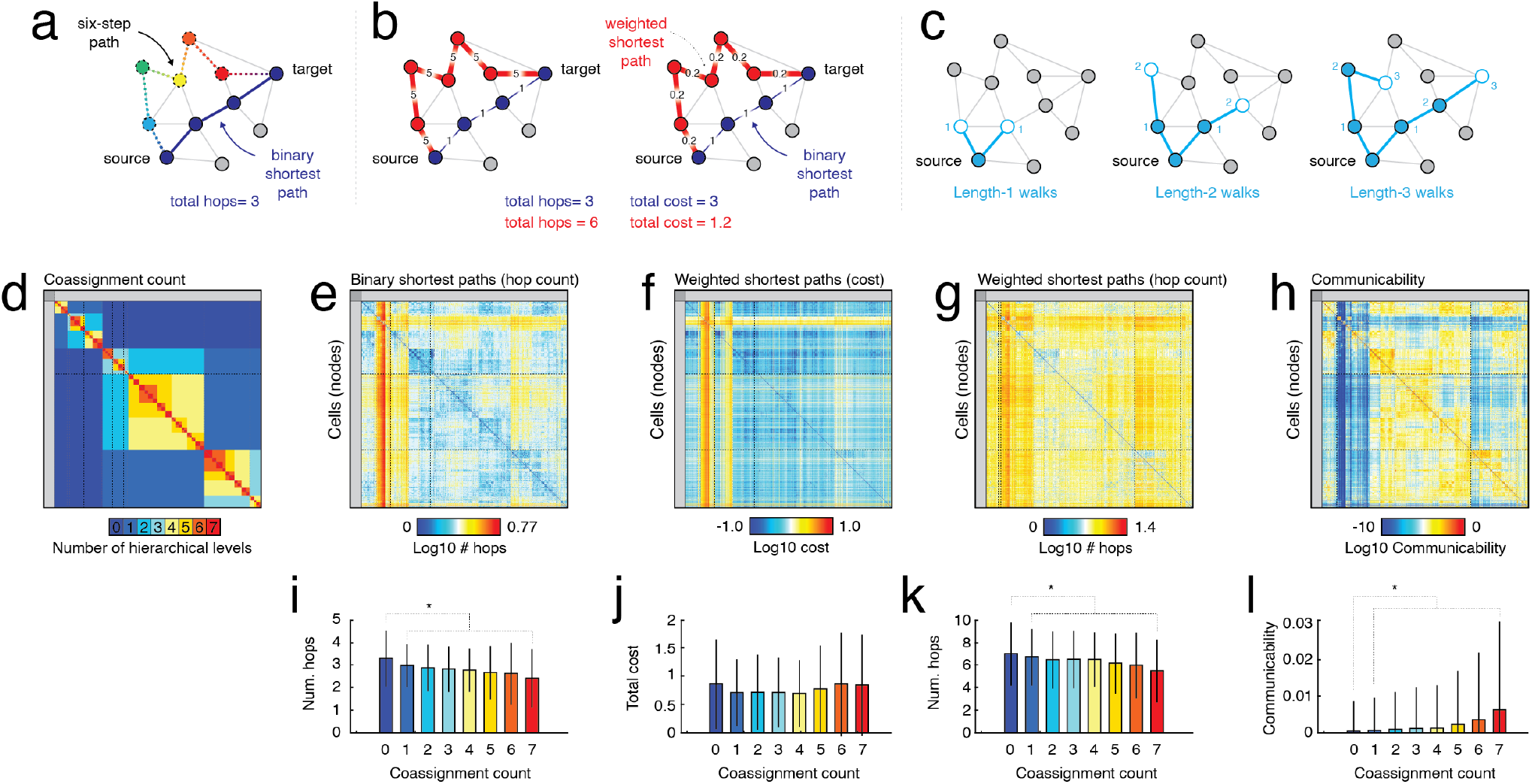
Linking processing paths to hierarchical community structure. (*a*) Schematic illustrating example path and shortest path between source and target nodes in a binary network. (*b*) Example of shortest weighted path. Note that weights are typically transformed from measures of affinity to measures of cost in the estimation of shortest weighted paths. Note also that the shortest weighted path may include more hops than the shortest binary path. (*c*) Illustration of walks of different length. Communicability counts and sums the number walks of all lengths between pairs of nodes, exponentially discounting the contributions from longer walks. (*d*) Hierarchical community coassignment matrix. Elements range from 0 (pairs of nodes never assigned to the same community at any level of the hiearchy) to 1 (assigned to the same community at every level). Panels *e*-*h* depict the binary shortest path, weighted shortest path (cost), weighted shortest path (hop count), and communicability between all pairs of nodes. For binary and weighted shortest paths, we find mean, mode, and maximum shortest paths of 3.28, 3, and 15 and 5.97, 5, and 23 hops, respectively (note that for the weighted shortest path, the quantity to be is minimized is not hop length, but a measure of cost estimated from connections’ weights). Panels *i* -*l* compare mean hop count, weighted shortest path (cost and hop count), and communicability between nodes at different levels of the community hierarchy.

#### 2. Weighted shortest paths

the least costly path between pairs of neurons irrespective of the number of hops (cost defined as 1*/*Number of synapses (Fig. 6*b*), and

#### 3. Weighted and directed communicability

a measure of how easily pairs of nodes can communicate using walks of any length, but with longer walks exponentially discounted (Fig. 6*c*). Two nodes linked by many two- or three-step walks can have high communicability, even if they do not share a direct connection.

We calculated these measures for every pair of nodes, generating *N × N* network communication matrices, where *N* is equal to the number of neurons. For every pair of nodes we calculated the number of hierarchical levels in which they were assigned to the same community (Fig. 6*d*) and linked those values to the three aforementioned measures (we show those matrices in Fig. 6*e-h*).

In general, we found that if nodes were assigned to the same community at any level of the hierarchy, their shortest path length and communicability tended to be smaller than nodes that were never in the same community (t-test, *p <* 10^−15^; Fig. 6*i,k,l*). Note, however, that we did not find this effect when we examine the cost matrix (Fig. 6*j*). We also found that hop distance and communicability decrease and increase monotonically with hierarchical levels, respectively, such that nodes assigned to the same community across all hierarchical levels tend to be connected *via* fewer hops and their walk density (indexed by communicability) was larger compared to those that appeared in communities less frequently (t-test; *p <* 10^−15^; Fig. 6*i-l*).

Taken together these results recapitulate well-established links between network modules and communication in the larval *Drosophila* connectome. These findings set the stage for more detailed analyses in future studies.

### Linking shortest-path trajectories to community hierarchy

In the previous section we showed that community hierarchy is, in aggregate, related to measures of communication. Here, we investigate those relationships in greater detail, focusing on shortest path trajectories from source to target nodes, detailing edge usage relative to nodes’ positions in the community hierarchy.

Specifically, we calculated the module co-assignment matrix so that every pair of nodes was assigned a value between 0 and 7 depending on the number of hierarchical levels in which they were both assigned to the same module. We then “masked” the co-assignment matrix with the binarized connectivity matrix, setting to zero all entries in the coassignment matrix corresponding to pairs of nodes that were not directly connected (Fig 7a). Using this re-labeled matrix, we tracked how often and where edges with labels 0, 1, 2, 3, 4, 5, 6, and 7 appeared in shortest paths. Finally, for paths of all lengths, *L*, we calculated the typical trajectory with respect to edge labels, which reveals how shortest paths travel across modular hierarchies on their way from a source to a target cell (Fig. 7*b*). Note that this approach is similar to the model studied in Winding *et al*. [21], wherein “seed neurons” probabilistically activate their post-synaptic partners, creating “cascades” of activation that propagate across the connectome. Using that model, the authors characterized the timing of activations. Here, we focus on shortest path structure, which represents an extreme case of the cascade model, corresponding to a 100% probability of activating post-synaptic partners. A further distinction between shortest paths and the cascade model is that neurons in the cascade model can enter a “deactivated” state following its own activation, wherein the neuron cannot activate its postsynaptic neighbors.

**Figure 7.**
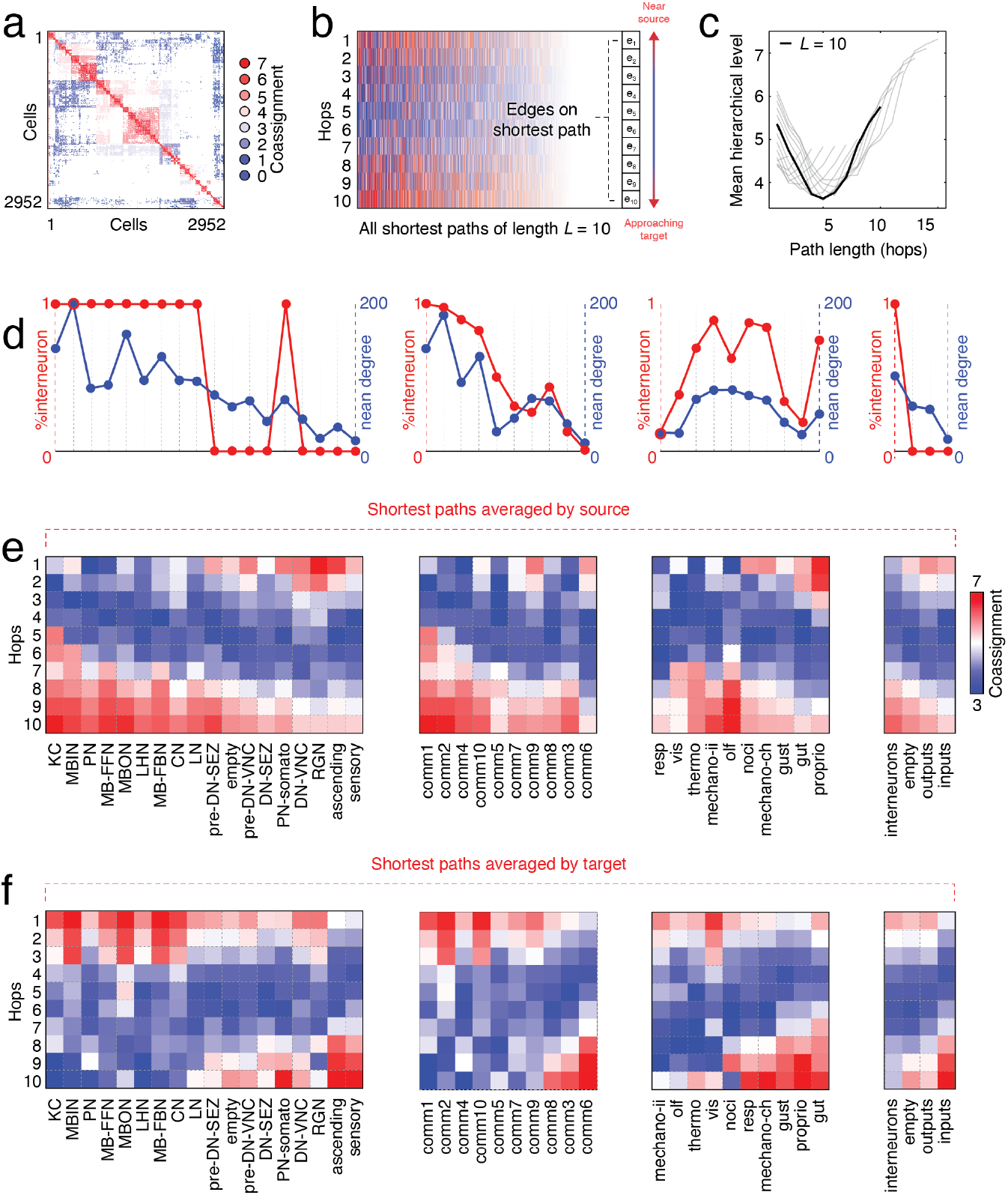
Shortest path trajectories. (*a*) Masked connectivity matrix. Edges are labeled based on coassignment probability: 0, 1, 2, 3, 4, and 5 indicate the number of hierarchical levels that a given pair of connected nodes were assigned to the same community. (*b*) We can then describe the composition of the shortest path from a given source node to a given target node in terms of the coassignment probabilities of the edges along that path. As an example, we show all shortest paths of length *L* = 10. (*c*) We then calculated the typical trajectory as the mean across all shortest paths, which revealed a characteristic u-shaped trajectory. That is, the first hops away from a source node tend to be to other nodes in the same community as the source node, followed by a series of cross-community hops, before terminating through a series of within-community hops as the shortest path approaches the target node. Panel *c* shows the characteristic trajectory averaged over all source/target pairs for *L* = 2 to *L* = 15 (there were very few shortest paths beyond this length and, for brevity, we ignore them). We then asked how these shortest path trajectories varied across cell types, communities, functional annotations, and macro cell groups. We found that, while most shortest paths follow a general u-shape, a subset deviate from this shape. For instance, shortest paths originating from Kenyon cells almost immediately begin using between-community connections to leave their local community, followed by a quick “ascension” along edges that fall within communities (see panel *d*). We find an opposite trajectory when we average only shortest paths that end with Kenyon cells (see panel *e*). These effects can be explained by two observations. Kenyon cells are interneurons and, like other interneurons, tend to be high degree. We find that groupings that deviate from the u-shaped trajectory are both dominated by interneurons and exhibit high average degree (see panel *c*).

We found that, irrespective of path length, shortest path trajectories tend to be initiated using edges that link to nodes in the same community as source node, advance to cross-community edges, before once again traveling along within-module connections near the vicinity of the target node. These trajectories give rise to a characteristic “u-shape”. However, these characteristic trajectories are estimated by averaging over all pairs of source and target nodes, collapsing across considerable variation. How much do trajectories vary when the source and target nodes are selected based on specific annotational properties? To address this question, we estimated typical trajectories when the source/target nodes have different cell types, communities, functional profiles, and macro cell groupings (Fig. 7*c-e*).

Interestingly, we found heterogeneity across source/target types. For instance, Kenyon cells (KC) – a class of interneuron – deviated considerably from the characteristic u-shaped trajectory. As sources, they followed an increasing and near-linear trajectory, so that the initial step in the shortest path, rather than linking to another node in the same module, immediately takes advantage of cross-community connections. The rest of the shortest path is comprised of an “ascent” towards the target, using edges that are increasingly likely to link nodes in the same community (Fig. 7*d*,*left*). We observe a similar, but opposite, trajectory for shortest paths where Kenyon cells are the target (Fig. 7*f*).

In contrast, neurons associated with sensation (sensory neurons in Fig. 7*d,e*) follow a trajectory opposite that of Kenyon cells. These cells, along with descending and ascending neurons in the ventral nerve cord, are enmeshed within segregated and assortative modules. As sources, the first steps on their shortest paths tend to be to other cells in the same group, eventually reaching cells that make cross-module connections (the trough of the u-shaped trajectory) and ascending *via* within-module connections towards the target neuron.

What explains the variation in shortest path trajectories? One hint comes from the trajectories of the macro cell groups. Interneurons–which include KCs– follow monotonically increasing and decreasing pathways when they are grouped by source and target, deviating from the characteristic u-shaped trajectory (Fig. 7*b*). Indeed, we find that the percentage of interneurons in each group, with the exception of the functional annotations, is largely in agreement with the ordering of trajectories, from those that are similar to the trajectories of Kenyon cells to those that are more u-shaped (Fig. 7*c*, red curve). This observation suggestions that the cell type composition of groups, namely the fraction of cells labeled as interneurons, helps determine the extent to which their respective trajectories deviate from the grand average.

What network property (properties) do interneurons exhibit that might explain this phenomenon? Relative to input and output units, interneurons tend to be high degree–i.e. they maintain relatively high numbers of incoming/outgoing connections. This aligns with the earlier observations that rich-clubs and core-periphery structures are enriched for interneurons. From the perspective of shortest paths, by virtue of being high-degree and non-assortative, interneurons can use inter-modular connections to rapidly hop from their own module to another (unlike input and output neurons, which form largely assortative structures and whose initial steps along shortest paths tend to be to other neurons in the same module).

As a final analysis, we assessed to what extent the u-shaped trajectories could be attributed to community structure alone, or if their shape and variation revealed something distinct about the organization of the connectome. To address this question, we sampled networks from the fitted stochastic blockmodel (100 repetitions). For each sampled network, we repeated the above analyses. We found that the the shortest paths of the sampled networks also exhibited u-shaped trajectories consistent with those previously reported (Fig. S14). Further, we also find that, given these null models, we recover the functional, cell type, group type, and community specificity of shortest paths.

Collectively, our findings suggest the u-shaped trajectories and their specificity are directly related to the nested community structure. That is, our null models preserve topological features of the network—community labels exactly and nodes’ in-/out-degrees approximately—but are otherwise not directly informed by neurons’ annotations. However, the results of our study suggest that many topological features of the larval fly connectome are also correlated with annotations; communities, rich clubs, and local connectivity properties (neurons’ degrees) are related to function/cell type/group type. Accordingly, in preserving these topological properties, we are likely inadvertently allowing information about annotations to “leak” into our analysis, further underscoring the need for the development of novel null models for disentangling connectivity data from annotations.

These observations, which are directly in line with previous large-scale connectome studies [53, 57, 61, 62], suggest that the connectional properties of interneurons relative to other cell types may endow them with the ability to rapidly transmit signals to other downstream modules while readily integrating signals from upstream sensory units.

## DISCUSSION

Here, we analyzed the connectome of the larval *Drosophila*. The focus of our analysis was on its community structure–i.e. divisions of the whole network into smaller sub-networks. We presented evidence that the network exhibited hierarchical communities and that these communities were largely assortative, with some notable exceptions. We also showed that the communities, which were defined based only on connectivity alone, delineated functional groups, cell types, and macro cell groupings from one another. We reported evidence of a rich-club, comprised largely of high-degree interneurons. Although rich-club members were distributed across all communities (at a coarse scale), they tended to concentrate within a small subset of communities. We showed the community boundaries were related to the ease of communication between neurons, and that in delivering a signal from one neuron to another, shortest paths traverse the hierarchy of communities in a specific way, but deviate when the source/target groups contain high percentages of high-degree interneurons.

### Communities reflect cell types and function

Community structure is one of the most studied properties of networks. It has been given proper mathematical treatment for at least half of a century, with early studies in sociology focusing on detecting communities using blockmodels fit to social network data [63–67]. The interest in discovering latent cluster structure in network data has continued to the present day [31, 32, 68], extending far beyond the social sciences, and, with new approaches and insights, community detection and analysis continues to be a quickly evolving sub-discipline within network science [69, 70].

In network neuroscience, especially, community structure has played a central role in shaping our understanding of brain network organization [17, 19, 71–75]. At the large scale–where most network neuroscience applications have occurred–communities are generally thought to reflect functionally related groups of neural elements [76] as evidenced by their circumscription of regions that activate under similar context [77, 78].

However, establishing direct links between modules and other annotational data–e.g. cytoarchitectonic properties and cell types–has been less successful at the large scale. In fact, only recently has a framework for curating, sharing, and comparing brainmaps become available [42]. Even in network science proper, there has been skepticism as to whether detected communities have any basis in “ground truth” community annotations, with studies reporting only modest alignment [70, 79].

Here, and in line with Winding *et al*. [21], we show that communities estimated based only on connectivity nonetheless are “enriched” for distinct modes of function, cell types, and cell groups. This means that cells carrying these annotations are concentrated within particular communities at a level unexpected by chance. Critically, we show that at coarse scales the enrichment patterns are nearly one-to-one, so that if a particular functional or cellular annotation is enriched in community “*X*”, it tends to not be enriched in other communities. These findings establish a correspondence between community structure and neurobiological annotation data, reifying the intuition and hypothesis that connectivity-defined communities reflect function and cytoarchitectural properties at the micro-scale.

An interesting extension of this work would be to incorporate the annotation data directly into the generative model and allowing variation in annotation data across neurons to interact and impact connection probabilities [80, 81]. Indeed, recent extensions of the stochastic blockmodel have made this possible by including annotations as a fixed set of parameters [82, 83]. To tease apart which annotations “drive” the formation of communities, one could systematically “lesion” different annotations from the set of parameters used to fit the blockmodel, assessing how the exclusion of different annotation types hinders goodness of fit measures.

### Communities are mostly assortative

Typically, when the term “modularity” is invoked in the context of biological neural networks, it is used to refer to “assortative” community structure – i.e. divisions of a network into segregated sub-networks. This type of structure is thought to support specialized function [84], promote evolvability [85], adaptability [86], robustness to perturbations [87], separation of dynamical timescales [88], and allow for efficient embedding of the network’s elements and wiring in three-dimensional space [14].

Indeed, empirical studies of brain networks have consistently revealed precisely this type of organization. On one hand, these observations could be viewed as evidence that, from the perspective of embodied nervous systems, assortative communities are functionally adaptive features for all (or just some) of the reasons mentioned earlier. On the other hand, the preponderance of assortative modules could also reflect biases in network construction – e.g. correlation-based metrics of functional coupling exhibit transitive relationships that might artificially reinforce assortative groupings [89] – or biases in the methods used to detect communities. The *de facto* community detection methods in network neuroscience – modularity maximization and Infomap – explicitly seek assortative communities. In other words, even if other types of community structure were present in a network, these methods are incapable of detecting it.

Stochastic blockmodels, though not without limitations [90], are capable of detecting generic community types, including assortative structure [29], and therefore offer a useful framework for assessing evidence of assortativity in brain networks. If there is a preponderance of statistical evidence supporting core-periphery of disassortative communities, then the stochastic blockmodel will recover said communities; on the other hand, if the evidence favors assortative structure, then the opposite will be true.

Here, we use a hierarchical variant of the stochastic blockmodel to detect communities. We find that, overwhelmingly, communities interact assortatively – i.e. most community pairs maintain stronger within-community connection density than between-community density. This observation is in line with other studies that have applied SBMs to connectome data [34, 35, 49]. Here, also in line with large-scale imaging studies [33], we find that the most assortative communities are associated with (pre) descending neurons involved in proprioceptive, nociceptive, and mechanosensory function – i.e. sensation and perception.

Although most community interactions are assortative, a small fraction (about 6%) are non-assortative. These motifs represent deviations from the near-uniform assortative interactions observed across the larval *Drosophila* nervous system and cross-linking communities to one another. These deviations also demand functional explanations; while assortative communities are thought to support specialized sensory processing, what are the functional roles of disassortativity and core-periphery structures? Future modeling studies should investigate these and related questions in greater detail. An interesting possibility not explored here, is that core-periphery structure emerges from overlapping assortative communities–i.e. if nodes are allowed to participate in multiple assortative communities, a group of nodes with co-membership to many of said communities can appear “core-like” [91]. In this case, the core-periphery structure observed here may be consistent with the hypothesis that neurons are organized into functionally specialized assortative communities, with the caveat that neurons may exhibit mixed memberships. However the hierarchical variant of the stochastic blockmodel studied here does not permit overlapping cluster structure. Future studies should investigate this phenomenon in greater detail.

### Hubness of interneurons gives them unique network properties

Heterogeneous and heavy-tailed degree distributions are a hallmark of real-world networks [92]. This distribution implies that, while most nodes make relatively few connections, a small and exclusive subset make disproportionately many connections, placing those nodes in highly influential positions within the network.

Indeed, some of the earliest network analyses of biological neural networks revealed the presence of high-degree and highly-central “hub” nodes [4], an observation that was further refined when it was discovered that hubs link together, forming “rich-club” structures [12]. These phenomena are hardly a product of networks reconstructed from magnetic resonance images, and have been observed across spatial scales and phylogeny [47, 93–96].

Here, we find that the neuronal connectome of the larval *Drosophila* also exhibits a heavy-tailed degree distribution and that high-degree neurons are comprised largely of interneurons situated in the mid-/central-brain. By virtue of making many connections, these neurons are conferred a number of unique structural (and possibly functional) properties. For instance, high-degree interneurons are the most likely to participate in non-assortative community motifs and rich-clubs. Their status as highly-connected units also positions them as key nodes along putative communication pathways (shortest paths), facilitating efficient and increasingly direct routes to downstream target neurons, while also making themselves easily accessible targets for pathways originating in other communities [61].

### Intra-community communication is more efficient than inter-community communication

Understanding how the configuration of a connectome’s edges shapes the flow of signaling has been a central aim of network neuroscience [97, 98]. Recent work has begun to address this question using network-based models of communication [57, 58] – stylized processes for delivering a “signal” from a source node to pre-specified target.

Here, we show that the effect of community structure is imprinted on the efficacy of communication processes – both centralized and decentralized. Specifically, we find that nodes consistently assigned to the same module across hierarchical layers are likely to be connected *via* shorter paths and exhibit greater “communicability” compared to nodes that are infrequently or never assigned.

Combined with the observation that communities are well-aligned with functional annotations and the observations made elsewhere that communication measures are strongly correlated with the magnitude of functional coupling between neural elements [55, 56], our findings position constraints imposed on communication by community structure as a key determinant of a neuron’s functional repertoire.

### Multi-scale network neuroscience

For the past two decades, network science has permeated virtually every scientific discipline [99]. Part of its success is owed to the generality of network models; a system’s details are abstracted away, leaving behind a set of circles (nodes) and lines (edges), representing those elements and their pairwise interactions.

This model has proven profoundly useful in neuroscience. In particular, it has been applied extensively to inter-areal connectome data [4–6, 100]. These analyses have identified core sets of phylogenetically conserved architectural features, the most commonly cited being small-worldness [10], hubs and rich clubs [4, 12], modular structure [101], and wiring cost reduction [39, 102, 103]. Though referring to static architectural properties, these features are often interpreted in terms of brain function; modules for specialized and segregated information processing, hubs and small-worlds for integration, and wiring cost as a constraint that limits the total material and metabolic expenses of the brain.

However, due in large part to the paucity of whole-brain, neuron-level connectome data, whether similar organizational principles are evident at the microscale remains unclear [104, 105]. As we firmly enter the era of nano-connectomics, it is becoming possible to not only assess whether features described in other scales are evident [106], but to understand altogether new network phenomena and uncover previously undetected properties [107]. Ultimately, this approach holds promise for effectively bridging scales. Starting from magnetic resonance imaging data, the smallest characterizable unit is the voxel or surface vertex; probing features at finer spatial scales is impossible. On the other hand, beginning with nano-scale data, we can coarse-grain our way to the scale of voxels (≈ 1 mm), presenting an opportunity for truly multi-scale network models [108].

### Future directions

The focus of this paper was to link network communities derived from synaptic connectivity with neuronal annotations. One of the challenges associated with this type of analysis is adequately addressing, from a statistical perspective, the nested nature of the detected communities as well as the annotations. For instance, we found that community 5 was significantly “enriched” for sensory neurons. That is, the community was comprised of more neurons with the “sensory” label than expected under the null model. However, we also found that community 5 was enriched for the labels “gustation” and “vision,” which are nested within the broader “sensory” label. To what extent should we anticipate this second result–enrichment for specific functional annotations– given that the same community was enriched for the “sensory” label? It is straightforward to construct counterexamples where the second outcome does not necessarily follow the first. For instance, a community could be enriched for “sensory” neurons but with each sub-category represented exactly proportional to its baseline rate. While there exist frameworks for dealing with nested hypotheses [109, 110], their application to large datasets with multiple levels of nestedness is not straightforward. With the proliferation of nano-scale connectome data [111] and increasingly rich and nested annotations of both neurons and their connections [112], this statistical issue presents a serious barrier. Future studies should focus on the exploration of frameworks for addressing this challenge.

A second important consideration for future studies concerns the enterprise of community detection and its role in network neuroscience. Community detection algorithms vary across multiple dimensions and, in general, will yield dissimilar estimates of community structure. We highlight an example of this here when we compare SBMs and modularity maximization; the SBM partitions suggest that communities are not strictly assortative, while modularity maximization is restricted to detecting assortative structure. We also compare our SBM partitions, which were derived using connectivity information alone, with partitions from Winding *et al*. [21], who used spatial information (hemisphere labels) in their clustering algorithm. In all cases, we found evidence for convergence across algorithms; communities were not identical, but had considerable overlap. Nonetheless, carrying out detailed comparisons of community detection algorithms may not be straight-forward in future studies; the increased dimensionality of nano-scale connectomes limits the application of computationally complex community detection algorithms. Relatedly, algorithms that incorporate meta-data and annotations run the risk of limited generalizability, i.e. they can only be applied to connectomes that have the same/similar meta-data. In general, these differences and considerations encourage the exploration and re-analysis of the same dataset using multiple approaches. Indeed, when viewed through the lens of statistical inference, there exists a many-to-one mapping of communities to connectomes [113], such that very different community structure can offer equally good/bad descriptions of a network depending upon the exact function that maps maps community labels to connectivity.

## MATERIALS AND METHODS

### Dataset

We analyzed the larval *Drosophila* connectome as published in [21]. The complete connectome included a giant strongly connected component of *N* = 2952 neurons and *M* = 110677 edges. Edges were weighted by synapse count for a total of *K* = 352611 synapses. The total network has a density of *δ* = 0.0127.

In Winding *et al*. [21], the authors analyzed these same data. In some cases they studied a thresholded version of the connectome, retaining only multi-synaptic connections. However, in others, the analyzed the complete and unthresholded connectome. Here, we applied no thresholding. Consequently, synapse counts ranged from 1 to 121. Note also that in Winding *et al*. [21], the authors distinguished between axon-to-axon, axon-to-dendrite, dendrite-to-axon, and dendrite-to-dendrite synapses. Here, we aggregated these four synapses classes together, opting to study the combined network.

We denote the presence/absence of a connection between pre-synaptic neuron *i* and post-synaptic neuron *j*–i.e. the directed adjacency relationship–as *A*_*ij*_; the number of synapses–i.e. the connection weight–is given by *W*_*ij*_.

### Hierarchical stochastic block model

Here, we fit a hierarchical stochastic blockmodel to the larval *Drosophila* connectome following [30] (https://graph-tool.skewed.de/). Stochastic blockmodels (or SBMs) are generative models of a network in that, given the community assignments of all nodes, *σ* = *{σ*_*i*_*}*, the model generates the observed network with probability: *P* (*W*|*θ, σ*), where *θ* refers to any additional model parameters that specify the link between community labels and the network. The hierarchical variant recursively fits the stochastic blockmodel to network data using an efficient agglomerative algorithm to minimize the posterior probability that the model generated the observed network [114].

We tested six variants of the hierarchical blockmodel. Specifically, we consider three different discrete distributions from which edge weights are drawn (“geometric”, “poisson”, and “binomial”). For each distribution type, we also consider a version of the model in which nodes’ degrees are treated as a set of parameters (the so-called “degree-corrected” model) and another that does not. In each case, we minimize the description length of the model and further refine the fit using Markov chain Monte Carlo procedure [115]. For each of the six models, we repeat this procedure with 100 random restarts.

To adjudicate between models, we compared description lengths (an information theoretic measure of how well each model compresses the observed network). We found that the description length of the degree-corrected model with a geometric distribution was smaller than that of all other models, suggesting that this model should be preferred (*p <* 10^−15^; Fig. S9).

Given this observation, we obtained an estimate of the consensus partition following Peixoto [38]. Briefly, this procedure involves initializing a degree-corrected blockmodel with a discrete geometric distribution over edge weights. We then equilibrate the Markov chain, and sample 10000 high-quality partitions. From these partitions, we align partitions to one another and estimate partition modes. This procedure identified a single mode, suggesting that the 10000 samples were likely noisy estimates one stable solution. We then estimated the “consensus” community assignment for each neuron. All analyses reported in the main text are carried out on this consensus solution.

### Hierarchical (recursive) modularity maximization

Modularity maximization defines communities to be groups of nodes whose observed density of connections maximally exceeds that of a null model. This intuition is formalized by the modularity heuristic:

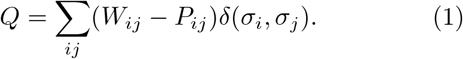

In this equation, *W*_*ij*_ is the observed weight of the connection between nodes *i* and *j, P*_*ij*_ is the expected weight of the same connection under a user-defined null connectivity model, *σ*_*i*_ is the community assignment of node *i*, and *δ*(*x, y*) is the Kronecker delta function, which evaluates to 1 when *x* = *y* and is 0 otherwise. As a null connectivity model, we specify the degree-preserving “configuration model”, under which 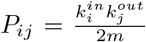 where 2*m* = ∑_*i*_ *k*_*i*_. In general, the aim of modularity maximization is to select the community assignments so that *Q* is as large as possible.

Here, we applied modularity maximization recursively to generate a hierarchy of assortative communities [116, 117]. Briefly, this algorithm began with the complete network. We optimized 1000 times with a generalization of the Louvain algorithm [118] and used consensus clustering to obtain a representative partition of the network [119]. For each of the *C* communities, we calculated its modularity contribution as *q*_*c*_ = ∑_*i*∈*c,j*∈*c*_(*W*_*ij*_ − *P*_*ij*_) and compared this value against a null distribution, generated by randomly permuting nodes’ community assignments. If the observed modularity contribution was unexpectedly large– i.e. *p <* 0.05, uncorrected–we retained the corresponding subgraph. Otherwise we discarded that community. Following this procedure, we obtain a series of subgraphs whose modularity contributions were bigger than expected; we propagated these subgraphs to the next hierarchical level, where we repeated the above procedure by detecting their modularity and testing whether the modularity contributions of the new sets of subgraphs exceeded their respective null distributions. This entire procedure was repeated indefinitely until no communities were propagated to the next level. At that point, the algorithm terminated, returning the (pruned) dendrogram of hierarchically related communities.

#### Community statistics

Let Γ_*c*_ be the set of all node indices assigned to community *c*. From this definition, we calculated a series of statistics.

1. *Community size* is defined as *N*_*c*_, or the number of elements in *c*.
2. The *synaptic density* of *c* is defined as 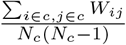.
3. The *binary density* of *c* is defined as 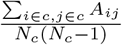 where *A*_*ij*_ is the binary adjacency matrix.
4. *Community diameter* is defined as max_*i*∈*c,j*∈*c*_ *D*_*ij*_, where *D*_*ij*_ is the inter-soma distance between nodes *i* and *j*.
5. *Mean distance* is defined as 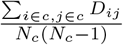.

### Coarse-graining and parcellations

In the supplementary material we show that synapse count (and its logarithm) are not correlated with inter-soma distance (Fig. S3). This was surprising, given that the so-called “exponential distance rule”, wherein the log of structural connection weights are approximately linearly correlated with measures of distance [39, 40, 103, 120].

However, most studies that described exponential-like distance rules analyzed inter-areal connectomes, wherein neurons are grouped into territories or areas (≈ 10^2^) and their synaptic connectivity averaged in some meaningful way. The resulting connectomes are, by definition, at a coarser scale than the single-cell connectome analyzed here.

To investigate whether coarse-graining the *Drosophila* connectome could recover an exponential-like distance rule, we partitioned neurons into clusters based on their soma *{x, y, z}* coordinates, resulting in spatially contiguous “parcels”. The weight of the connection between parcels *u* and *v* was calculated as the synapse count between all neurons in both clusters divided by the geometric mean of the two clusters’ sizes–i.e. 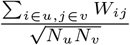. We also calculated centroids as the centers of mass for all neurons assigned to each cluster. From these centroids, we calculated an inter-areal distance matrix. Using these areal measures of connection weights and distances, we could calculate the correlation between the logarithm of connection weight and distance while varying the number of clusters, *k*. When *k* = *N*_*neurons*_, we recover single-cell resolution.

We also calculated a second measure of distance using neurons’ arbors. Each neuron’s arborization can be modeled as a tree network of connected nodes (with soma as parent). For a pair of synaptically coupled neurons, we can calculate the shortest path through these arbors from the pre-synaptic neuron to the synapse and from the synapse to the post-synaptic neuron. These paths are associated with two measures of distance: the first is the number of “hops” in the shortest path; the second is the Euclidean distane along the shortest path. That is, because the nodes of the arbor network are embedded in three-dimensional space, we can calculate the length of each segment and add up the total length of the segments that make up the shortest path. We focus on this second measure of distance–the shortest path Euclidean distance–which can be interpreted as the curvilinear length

We then randomly sampled 10000 values of *k*, independently partitioning neurons into clusters for each sample. For each sample, we calculated the correlation between the logarithm of connection weight and distance and the slope of that linear relationship.

### Community enrichment analysis

Both the stochastic blockmodel (main text) and modularity maximization (supplementary materials) partition neurons into non-overlapping groups. In parallel, the *Drosophila* connectome dataset was accompanied by a series of neuron-level annotations–e.g. cell type, function, and cell group. A natural question to ask is whether a community overlaps with any of these annotations.

To address this question we used an “enrichment” analysis. Intuitively, this approach calculates whether a particular annotation is concentrated within nodes assigned to a given community more than expected by chance. As an example, we show in the main text the case where we compare community six (from the second hierarchical level) with the *mechano-ch* functional labels (Fig. 3)*a,e*). We calculate the amount of overlap between these two clusters – i.e. the number of neurons that belong to both community six and the *mechano-ch* labels – and compare this count to a null model in which neuron locations are permuted while preserving, coarsely, their spatial statistics [44] (https://github.com/murraylab/brainsmash). We repeat the permutation process 1000 times and calculate the fraction of those 1000 iterations in which the overlap score was at least as large as the observed overlap. If this fraction (*p*-value) is small, then we conclude that community six is “enriched” for *mechano-ch* (borrowing the term “enrichment” from gene ontology studies). Note that in the main text, we correct for multiple comparisons using the procedure outlined by Benjamini and Hochberg [121], wherein the user specifies an allowable false positive rate and the critical value is adjusted to ensure that this rate is not exceeded. In all cases, we set the accepted false positive rate to 0.01 (1%). We pooled *p*-values across enrichment tests for function, cell-type, and group, and adjusted the adjusted *p*-value accordingly, resulting in a single statistical threshold that was applied to all tests. In Fig. S13 we explore an alternative and more conservative strategy for multiple comparison corrections. Namely, we use the Bonferroni correction, wherein our corrected critical value is penalized based on the number of tests we performed. As with the Benjamini and Hochberg [121] method, we penalize based on the total number of tests performed across functional, cell-type, and group-type annotations.

Note that this procedure can also be carried out with continuous variables. Later, we assess whether cell types, functional labels, and cell groups are enriched for community motif measures (assortativity, coreness, periphery, disassortativity, and the diversity index). As a concrete example, consider an effort to link the *interneuron* macro cell group with “coreness”. To do this, we calculate the mean coreness of all neurons labeled as interneurons. We then compare this number against a null distribution generated by permuting the spatial co-ordinates of the “coreness” measure, once again approximately preserving the spatial statistics of this whole-brain map.

### Non-assortative motifs

Unlike modularity maximization, which seeks assortative (cohesive) communities, the stochastic block-model is capable of detecting general community interactions, including core-periphery and dis-assortative structure. To detect these types of structures, we borrowed a technique from Betzel *et al*. [33, 34], that focused on pairwise “community motifs” (https://github.com/brain-networks/wsbm_sampler). Briefly, for a given pair of communities *r* and *s*, we calculated the [2 *×* 2] synaptic density matrix, *ω*, whose elements *ω*_*rr*_, *ω*_*ss*_, *ω*_*rs*_, and *ω*_*sr*_ represented the synaptic densities between nodes in *r* and *s*.

Given *ω*, we can define three types of motif interactions:

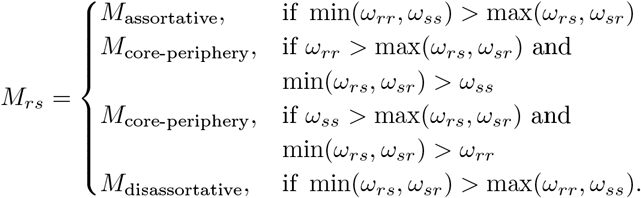

From these classifications we were able to associate motifs to individual nodes. Node *i*’s participation in motif *M* was calculated as the number of times that the community to which node *i* was assigned interacted with any other community to form a motif of type *M*. We then normalized these counts by the total number of motifs (for a *K*-community partition there are in total *K*(*K* − 1) total motifs). Importantly, when computing participation in core-periphery motifs, we distinguished between the core and periphery, and computed separate participation scores for each. Finally, from participation types we computed each node’s *diversity index*, which measured the entropy of its normalized participation in each motif type.

### Rich-club analysis

We also performed a rich-club analysis in the main text. Intuitively, rich-clubs are groups of hubs that are also highly connected to one another. This intuition is formalized by the rich-club coefficient, *ϕ*(*k*), which measures the density of connections among nodes whose degree is greater than *k* [122]. In weighted and directed networks, there are a number of decisions to make when calculating *ϕ*(*k*) [123]. Namely, how does one factor in connection weight and directedness? Here, we ignore weight, focusing on the binary rich-club. We also calculate *ϕ*(*k*) using nodes’ total degree–i.e. the sum of their incoming and outgoing connections.

The precise value of *ϕ*(*k*) is not particularly meaningful in isolation. Rather, it must be contextualized by comparing against a null model. Here, we generate null networks with identical degree sequence (incoming and outgoing) as the original network [50]. For each network, we also calculated the rich-club coefficient at every value of *k*. We then compared the observed coefficients with the null distribution, estimating a *p*-value for every *k*.

### Communication measures

Connectomes facilitate communication among neural elements. Therefore, connectome architecture plays a crucial role in shaping communication patterns [53, 54]. Here, we estimated communication capacity between every pair of nodes, *i* and *j*, based on three putative communication policies:

#### 1. Communication via binary shortest paths

This measure corresponds to the communication policy in which signals are selectively routed from one neuron to another *via* the shortest path – a sequence of nodes starting with *i* and ending with *j*, where every successive node corresponds to one of the outgoing connections of its parent, and whose total length (number of hops) is minimized.

#### 2. Communication via weighted shortest paths

This measure is similar to binary shortest paths, but assumes that the measure to be minimized is not hop count, but the cost of the shortest path. That is, each edge is assigned a cost – here, the reciprocal of its edge weight, i.e. *cost*_*ij*_ = 1*/W*_*ij*_ – and path with the shortest possible weight is detected algorithmically. Note that this communication policy allows us to estimate two measures of communication capacity for each pair of nodes, *i* and *j* – the total cost of the weighted shortest path as well as the total number of steps on that path.

#### 3. Communication via communicability

This measure calculates the number of walks of length 1, 2, 3, …, between nodes *i* and *j*, while exponentially discounting longer walks. Thus, two nodes that are not directly connected can nonetheless exhibit high levels of communicability provided that they are linked by many multi-step walks [124, 125]. Note that typically the communicability matrix is calculated as *e*^*W*^, where *W* is connectivity matrix and *e* is the matrix exponential. Here, however, we follow Crofts and Higham [124] and calculate a normalized communicability matrix that controls for nodes’ in-/out-degree (see **Materials and Methods**).

## Data availability statement

*Drosophila* connectome data used in the present study is available here: https://github.com/brain-networks/larval-drosophila-connectome

## Code availability statement

1. Code for estimating the hierarchical stochastic blockmodels is available here: https://graph-tool.skewed.de/
2. Code for estimating community motifs is available here: https://github.com/brain-networks/wsbm_sampler
3. Code for performing spatially-constrained permutations of brain maps: https://github.com/murraylab/brainsmash

---

**Figure S1.**
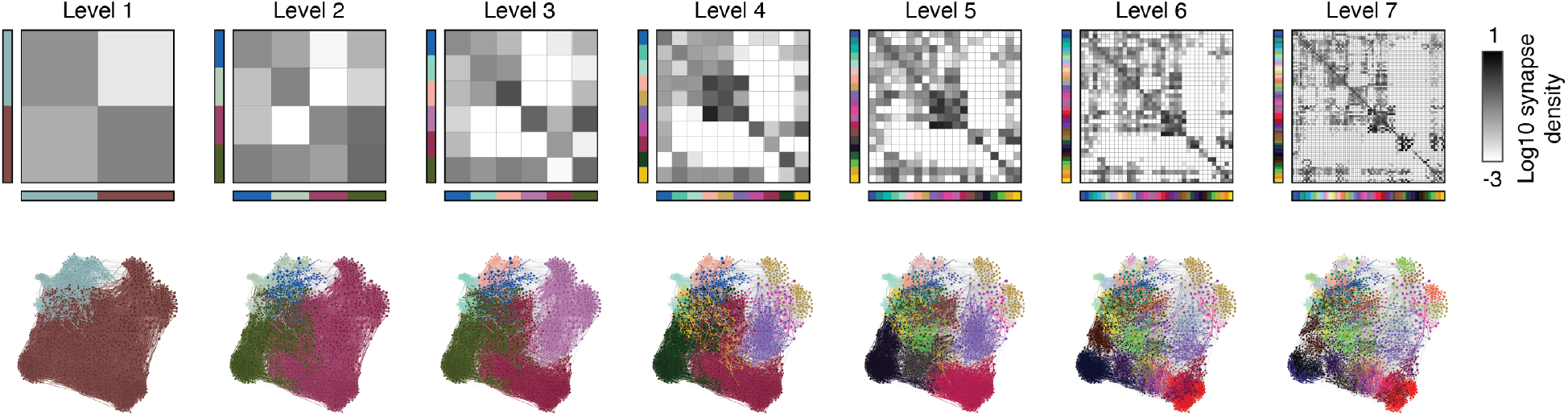
Coarse-graining the larval *Drosophila* connectome by community. Panels correspond to community-by-community matrices at each of the five hierarchical levels. The elements of the matrices correspond to synaptic densities – number of synapses divided by area of community-by-community block. The bottom row shows a reduced network (*N* = 2952 neurons and top six outgoing connections per neuron), with neurons labeled by community. Edges are colored if they fall within communities and left grayscale if they fall between.

**Figure S2.**
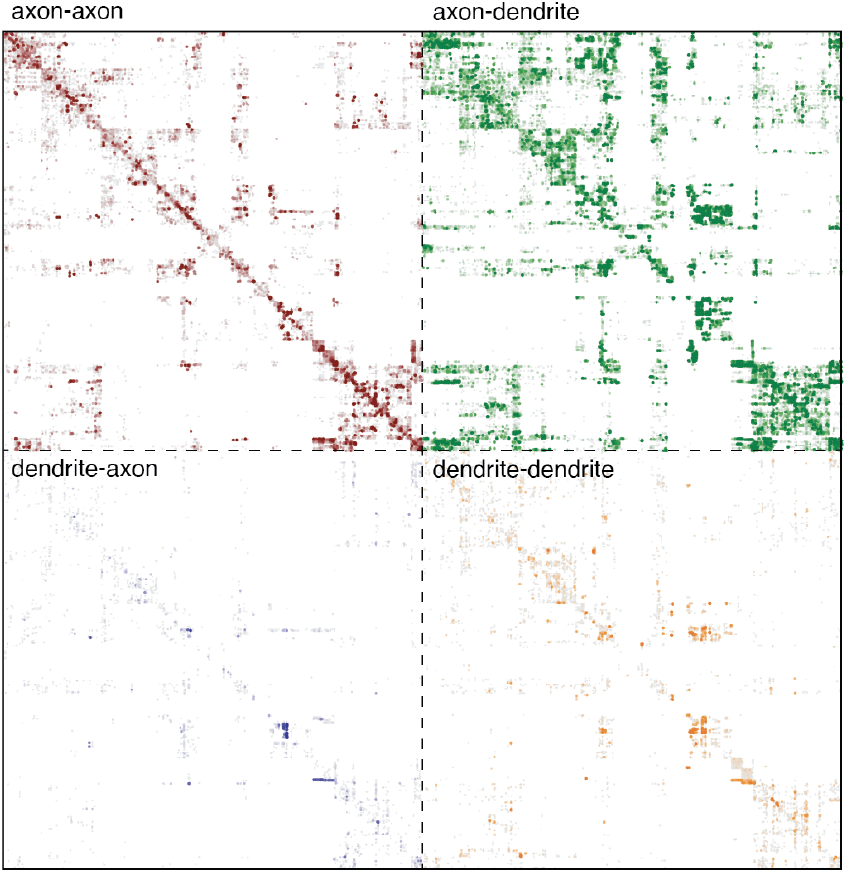
Connection-type matrices ordered by community label. In Winding *et al*. [21] the authors distinguish between axon-axon, axon-dendrite, dendrite-axon, and dendrite-dendrite synapses. Here, we show these matrices together. Note that the ordering of rows/columns for each matrix follows the ordering in Fig. 1.

**Figure S3.**
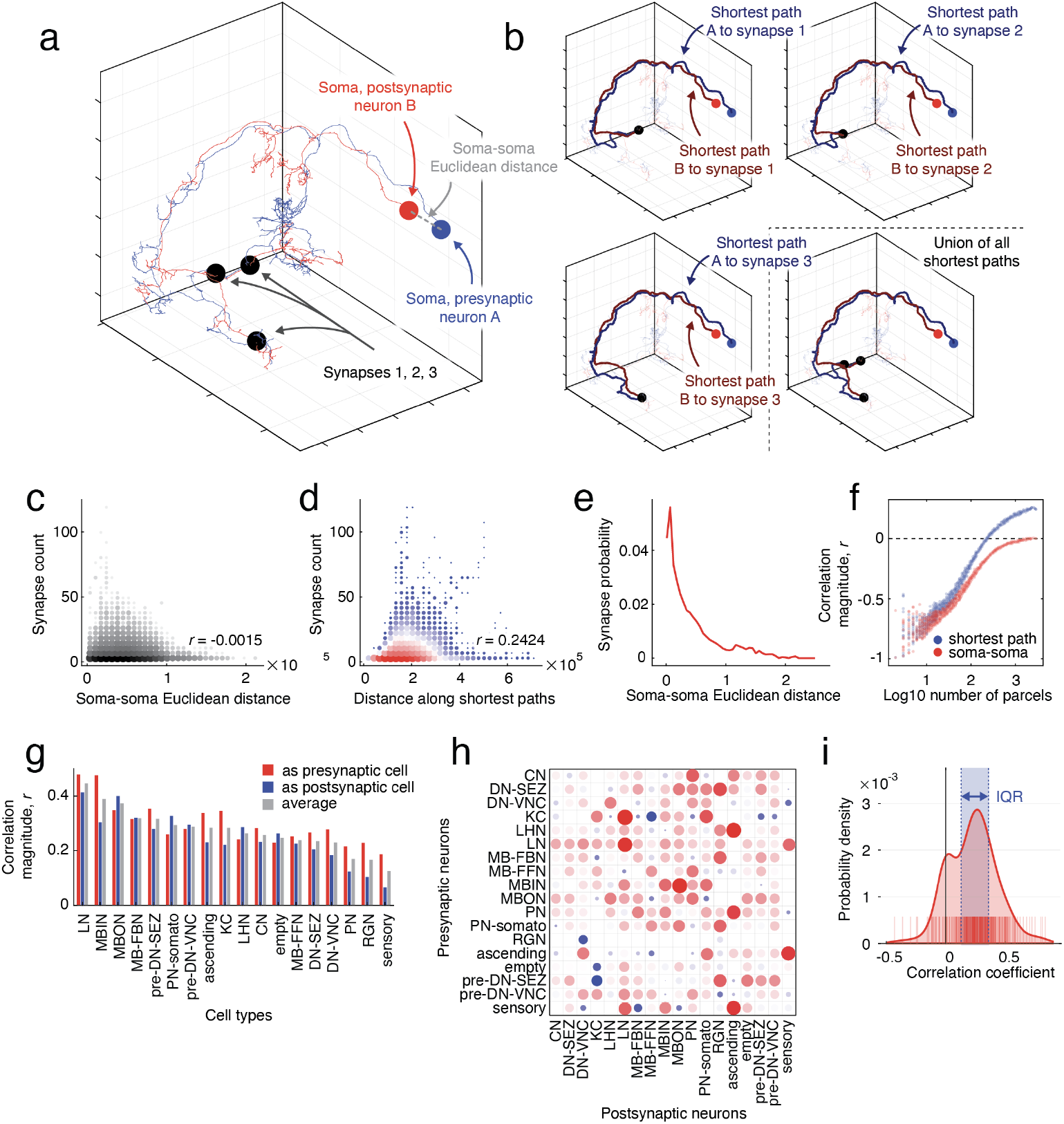
Assessing spatial dependencies at varying scales. Here, we analyze the spatial relationships between neurons. We focus on two ways of establishing a distance between pairs of cells. (*a*) The first is the straight-line distance between their somata (“soma-soma Euclidean distance”). We also calculate a second measure of distance based on the distance through the respective arbors of pre-/post-synaptic neurons to their synapses (shortest path from pre-synaptic neuron to synapse + shortest path from synapse to post-synaptic neuron; segments along the shortest paths are weighted by their Euclidean distance). For neurons connected by multiple synapses, we identify the “shortest paths backbone” – the set of node-to-node arbor segments appear in at least one shortest path – and calculate the distance as the total length (Euclidean distance) of segments in this backbone. We compared both of these metrics to total synapse connect. (*c*) Soma-soma Euclidean distance was not related to synapse count; (*d*) Shortest path distance was positively correlated with synapse count (*r* = 0.24; *p <* 10^−15^). (*e*) Nonetheless, as Euclidean distance between somata increasd, the likelihood of two neurons being connected by even one synapse decreases monotonically. Next, we asked how these relationships varied across different levels of coarse-graining. To address this question, we partitioned cells into spatially compact clusters using the *k*-means algorithm and soma-soma distance as the metric (randomly sampled *k* ∈ [2, 2952]) with 10000 random restarts). (*f*) We found that as the network gets coarse-grained, the we recover anticorrelations between the cluster-cluster weight and the separation between clusters (for both soma-soma and shortest paths distance). Next, we investigated how shortest paths distance varied for different cell types. (*g*) First, for each of the 18 cell types, we calculated the correlation between the shortest path distance to and synapse count of all postsynaptic partners (red). Next, we calculated the correlation between shortest path distance to and synapse count of all presynaptic partners (blue). Finally, we combined pre-/post-synaptic neighbors to calculate the total correlation (grey). We recapitulated the main global findings (panel *d*), reporting only positive associations. We also identified considerable heterogeneity across cell types, with local neurons (LNs) exhibiting the strongest association. (*h*) Finally, we investigated in even greater detail how the association between synapse count and shortest path distance varied for combinations of cells, e.g. connections from “local neurons” to “Kenyon cells”. Again, we found that nearly all correlations were positive (see panel *i*) but with considerable heterogeneity in terms of the strength of the association. The strongest associations were “KC-to-LC” (*r* = 0.84), “MBIN-to-MBON” (*r* = 0.81), and “LN-to-LN” (*r* = 0.75).

**Figure S4.**
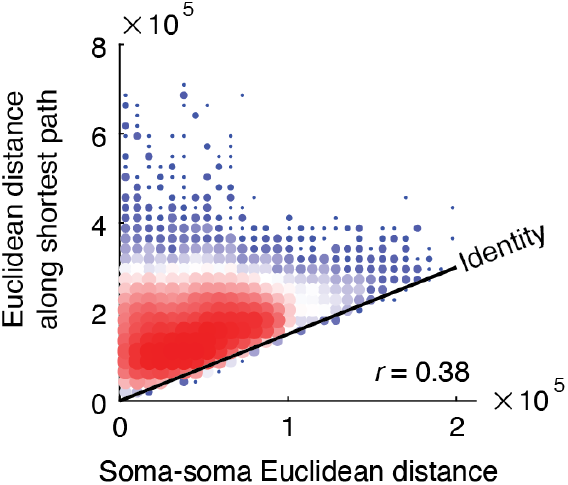
Direct comparison of soma-soma Euclidean distance with distance along s. We calculated two measures of neuron-to-neuron distance. The first was the soma-to-soma Euclidean distance. However, this measure does not account for the arborization of neurons and the locations of synapses (which are frequently far from cell bodies). We calculated a second measure of distance based on the path structure of pre-/post-synaptic neurons’ arbors. Here, we compare the two measures to one another as a 2-dimensional histogram.

**Figure S5.**
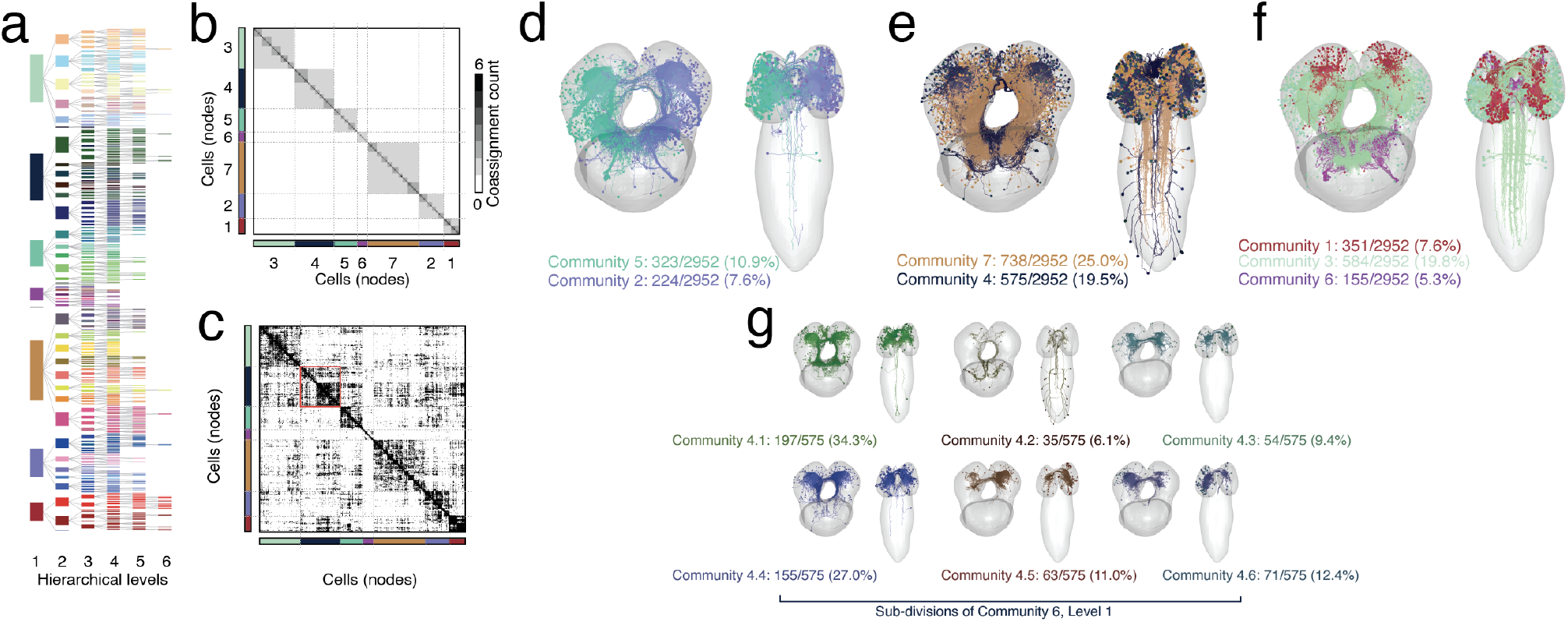
Example hierarchical communities obtained using modularity maximization. In the main text, we described communities detected a weighted and hierarchical stochastic blockmodel. Here, we show analogous figures for a hierarchy obtained using a recursive and hierarchical variant of modularity maximization. Note that communities here are assortative. (*a*) Dendrogram of communities. (*b*) Module co-assignment matrix. (*c*) Connectivity matrix ordered by communities. Panels *d, e*, and *f* depict level 1 communities. Note that, unlike communities obtained by the stochastic blockmodel, we find evidence of communities isolated within each hemisphere (see panel *d*). Panel *g* shows six of the communities obtained in level 2 after subdividing community 6 from the previous level.

**Figure S6.**
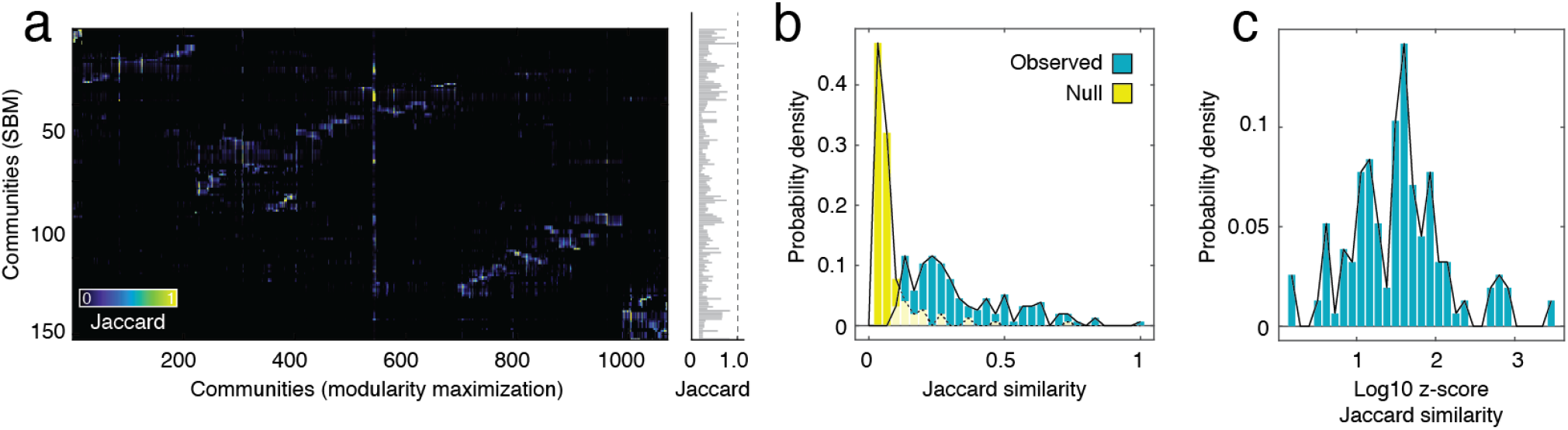
Correspondence between communities obtained with modularity maximization and the stochastic blockmodel. The hierarchical blockmodel and modularity maximization generate 155 and 1070 distinct communities, respectively. Here, we seek a correspondence between communities. Specifically, we calculate the Jaccard index between every pair of communities (see panel *a*, left). For each SBM community, we calculate the max correspondence with any community obtained with modularity maximization (panel *a*, right). (*b*) We compared, as a distribution, the maximum similarity scores from the observed data to the null distribution estimated by permuting community assignments. (*c*) We also z-scored each community’s maximum Jaccard similarity. Here, we show the logarithm of those z-scores (a score of “2” indicates that the observed Jaccard similarity was 10^2^ standard deviations greater than the mean of the null distributions).

**Figure S7.**
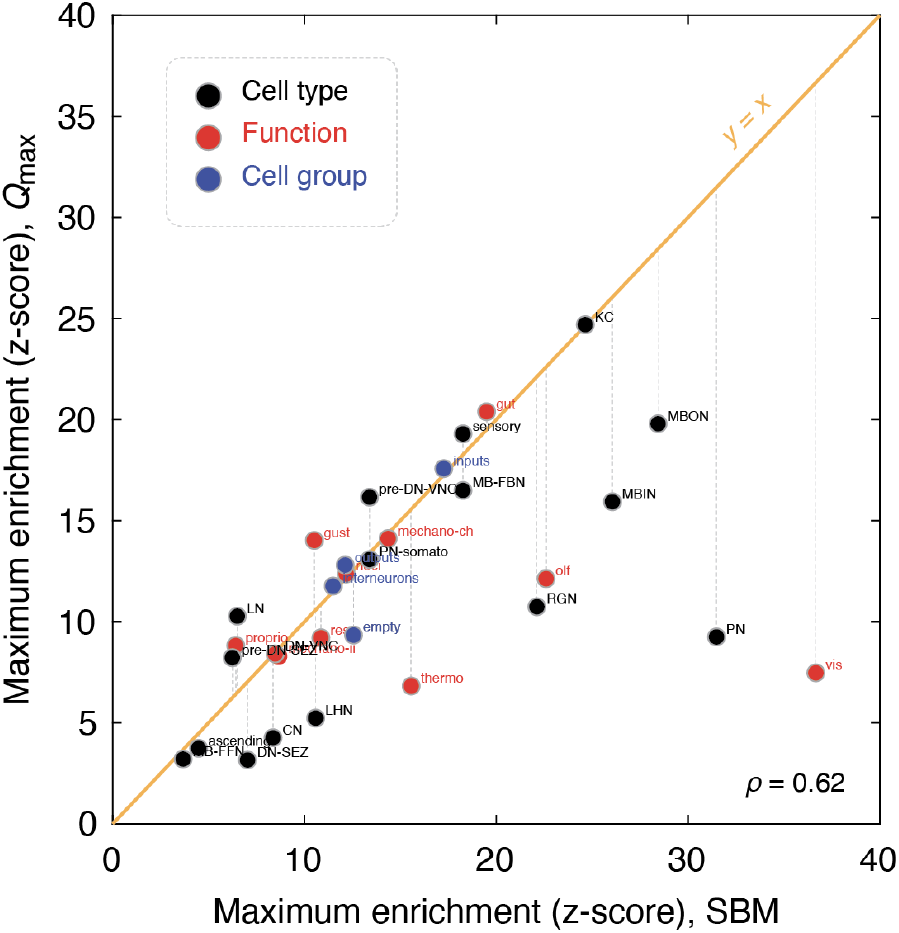
Comparing enrichment scores for blockmodel with modularity maximization. In the main text we calculated enrichment scores for annotation data relative to eight level-two communities. In the supplement we also describe communities estimated with modularity maximization. Using those communities, we repeated the enrichment analysis and, for each annotation, calculated its maximum enrichment score. Here, we plot the maximum enrichment scores obtained using the stochastic blockmodel with those obtained using modularity maximization.

**Figure S8.**
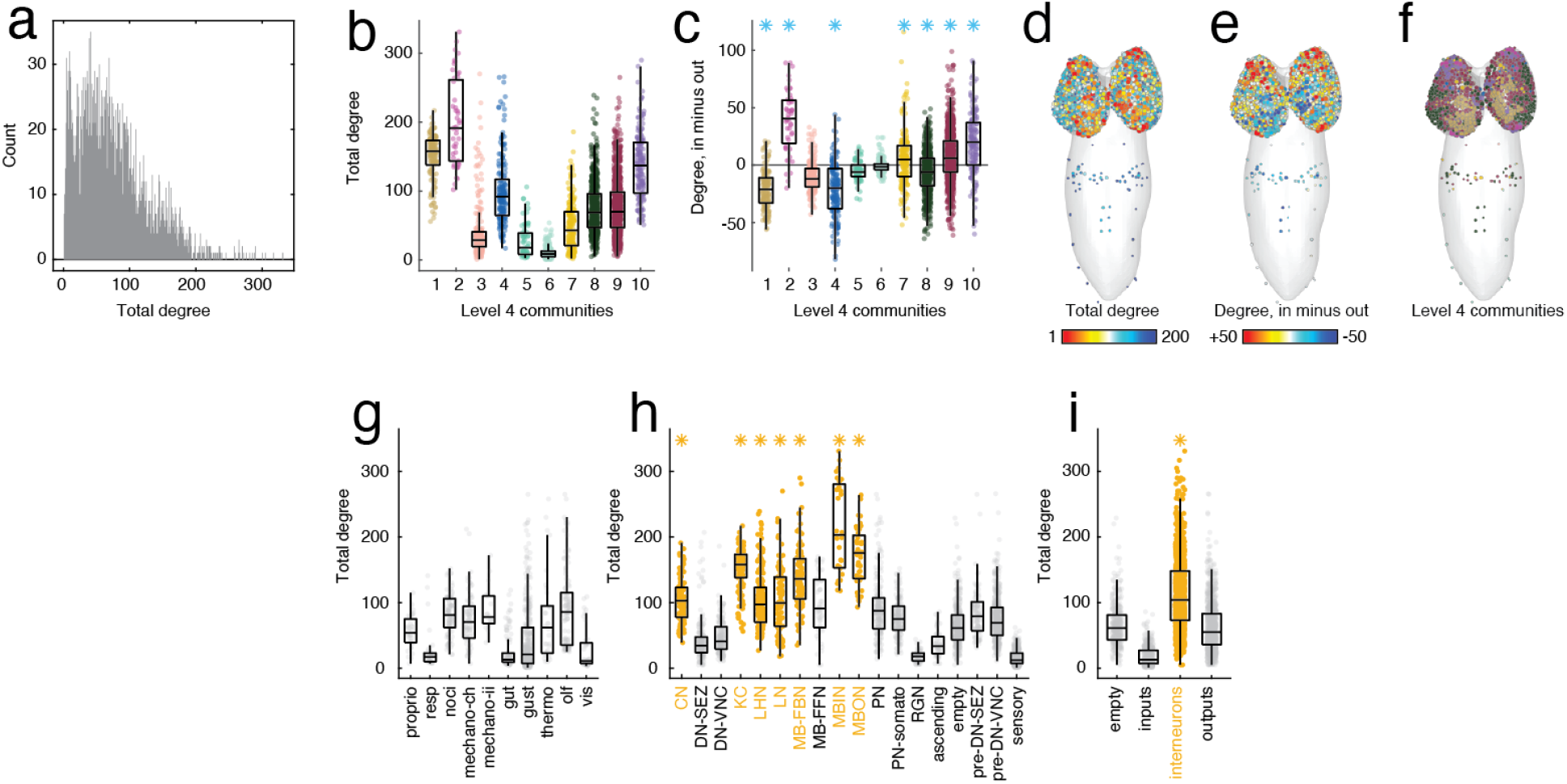
Analysis of high-degree hubs. (*a*) Degree distribution (combined in-/out-degree). (*b*) Total degree grouped by level two communities. (*c*) Differences in incoming *versus* outgoing degree, grouped by level-four communities. Panels *d, e*, and *f* show total degree, degree difference, and level two communities in anatomical space. Panels *g, h*, and *i* show degree grouped by functional group, cell type, and macro cell label. Yellow annotations are statistically significant; gray annotations are not.

**Figure S9.**
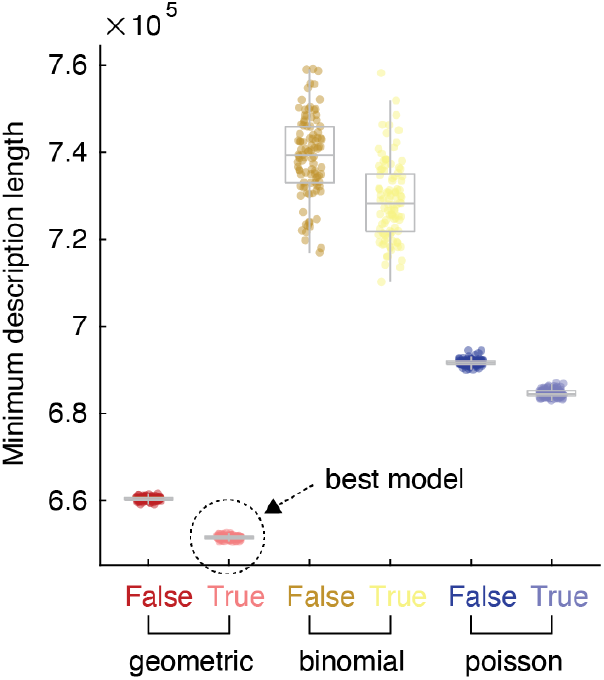
Description length. Comparison of minimum description length across six different nested stochastic block-models (each with 100 random initializations). We considered three distributions from which edge weights were drawn: “geometric”, “binomial”, and “poisson”. For each distribution type, we considered both degree-corrected (True) and non-degree-corrected (False) versions. Note that smaller description lengths are favorable.

**Figure S10.**
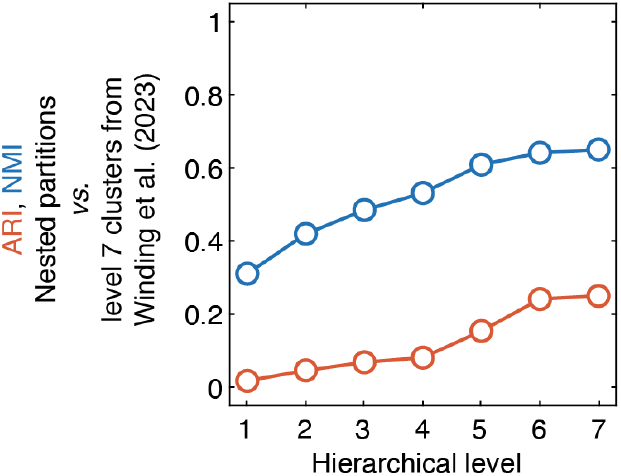
Similarity of nested communities from hierarchical blockmodel with those reported in Winding *et al*. [21]. In the supplement to the paper by Winding *et al*. [21], the authors share cluster labels for neurons (the 7th level of their own hierarchical model). We compare those labels with those detected using the nested/hierarchical stochastic blockmodel reported here in the main test. Specifically, we calculate the adjusted Rand index (ARI) and normalized mutual information (NMI) of each hierarchical level with the partitions shared by Winding *et al*. [21]. Both measures are bounded to the interval [0, 1] with larger values corresponding to greater correspondence between partitions. We make two primary observations. First, neither measure is close to zero. That observation suggests that the clusters reported here are not entirely dissimilar from those reported in [21]. Secondly, and on the other hand, neither measure approaches a value of 1, indicating that the partitions reported here are also not identical to those in [21]. In short, these observations suggest that the clusters in Winding *et al*. [21] and those reported here may capture distinct meso-scale patterns in the connectome.

**Figure S11.**
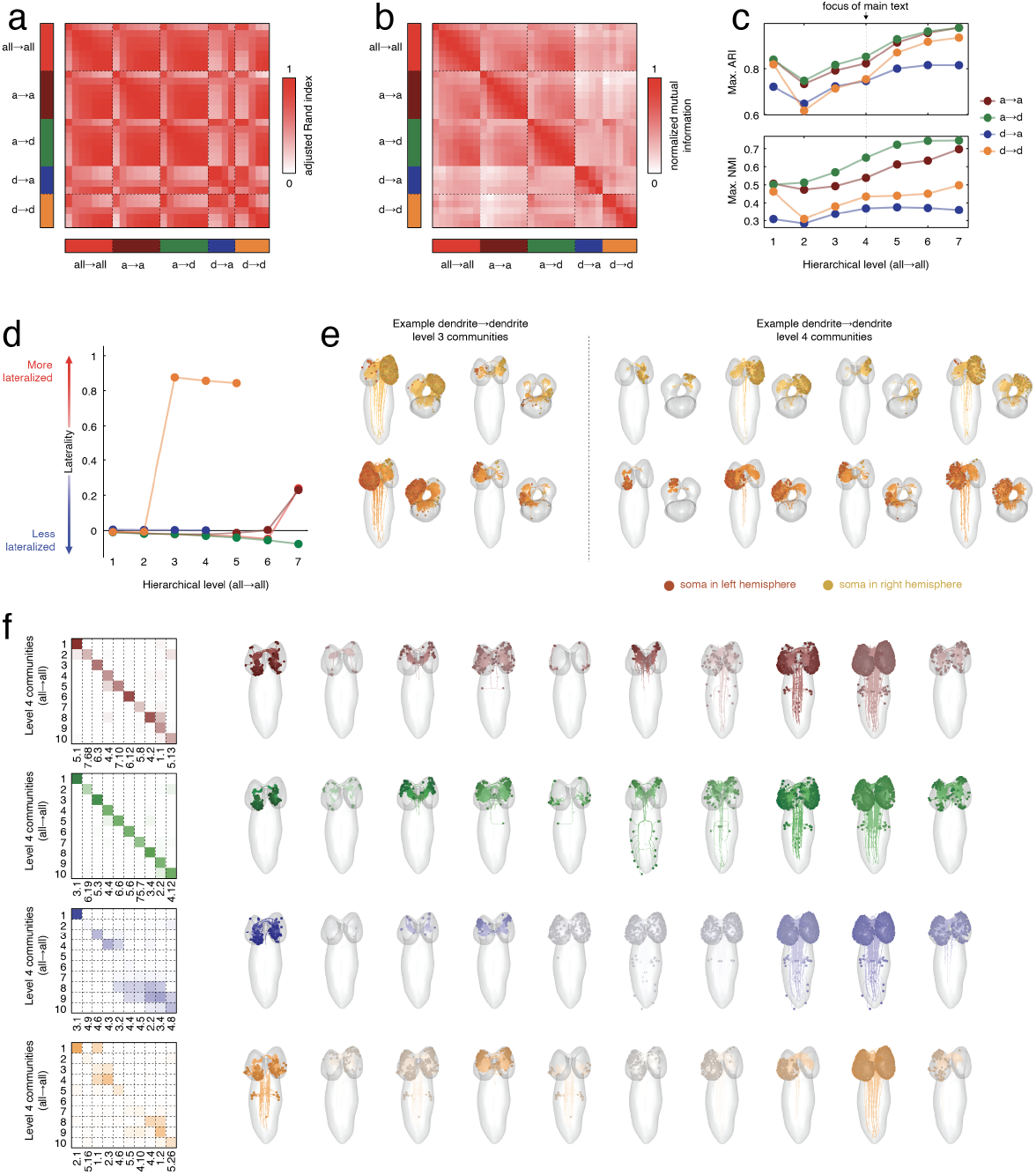
Synapse-type-specific community structure. In the main text we described the community structure of the *Drosophila* larval connectome. That network distinguishes between four synapse types: axon→axon, axon→dendrite, dendrite→axon, and dendrite→dendrite. However, in the main text we combined synapse counts to obtain a single connectivity matrix that summarized the total number of synapses between all pairs of neurons. Here, we repeat the community detection procedure separately for each of the four synapse-type-specific networks. Note that not all neurons make synapses of each type. Consequently, the networks whose communities we detected contained different numbers of nodes (2887, 2880, 1907, and 2204 neurons for axon→axon, axon→dendrite, dendrite→axon, and dendrite→dendrite networks, respectively). We found that the optimal nested partitions contained 7, 7, 4, and 5 hierarchical levels. Panels *a* and *b* compare levels of nested partitions to one another and to the combined (all→all) network using the adjusted Rand index and normalized mutual information. Larger values indicate greater levels of similarity. (*c*) For each hierarchical level of the all→all network communities, we identified the maximum similarity to *any* of the hierarchical levels detected in the synapse-type-specific networks. We find that axon→axon and axon→dendrite networks were most similar to the all→all network across its seven hierarchical levels. This likely reflects the fact that most synapses are either axon→axon and axon→dendrite and, consequently, are disproportionately represented in the all→all network. Next, we investigated why the dendrite→axon and dendrite→dendrite communities are so dissimilar. (*d*) Interestingly, we found that the dendrite→dendrite network exhibited highly lateralized communities [126]. That is, communities were comprised of neurons from either the left or right hemisphere, but not both. (*e*) Examples of lateralized communities for which there was a clear partner in the contralateral hemisphere. Finally, we asked how well the level 4 communities described in the main text were matched (at any hierarchical level) in the synapse-type-specific networks. We calculated the Jaccard index between each of the ten communities with every synapse-specific community. We then used the Hungarian algorithm [127] to optimally match all→all communities with synapse-specific communities. The heat maps on the left of panel *f* show the magnitude of correspondence (bound to the interval [0, 1]. White values are close to 0 and saturated colors are close to 1. The labels on the x axis indicate, first, the hierarchical level from which the optimal match originated and its label within that level. For example, a value of “3.2” indicates community “2” in hierarchical level “3”. For reference, we show to the right the topography of the optimal communities.

**Figure S12.**
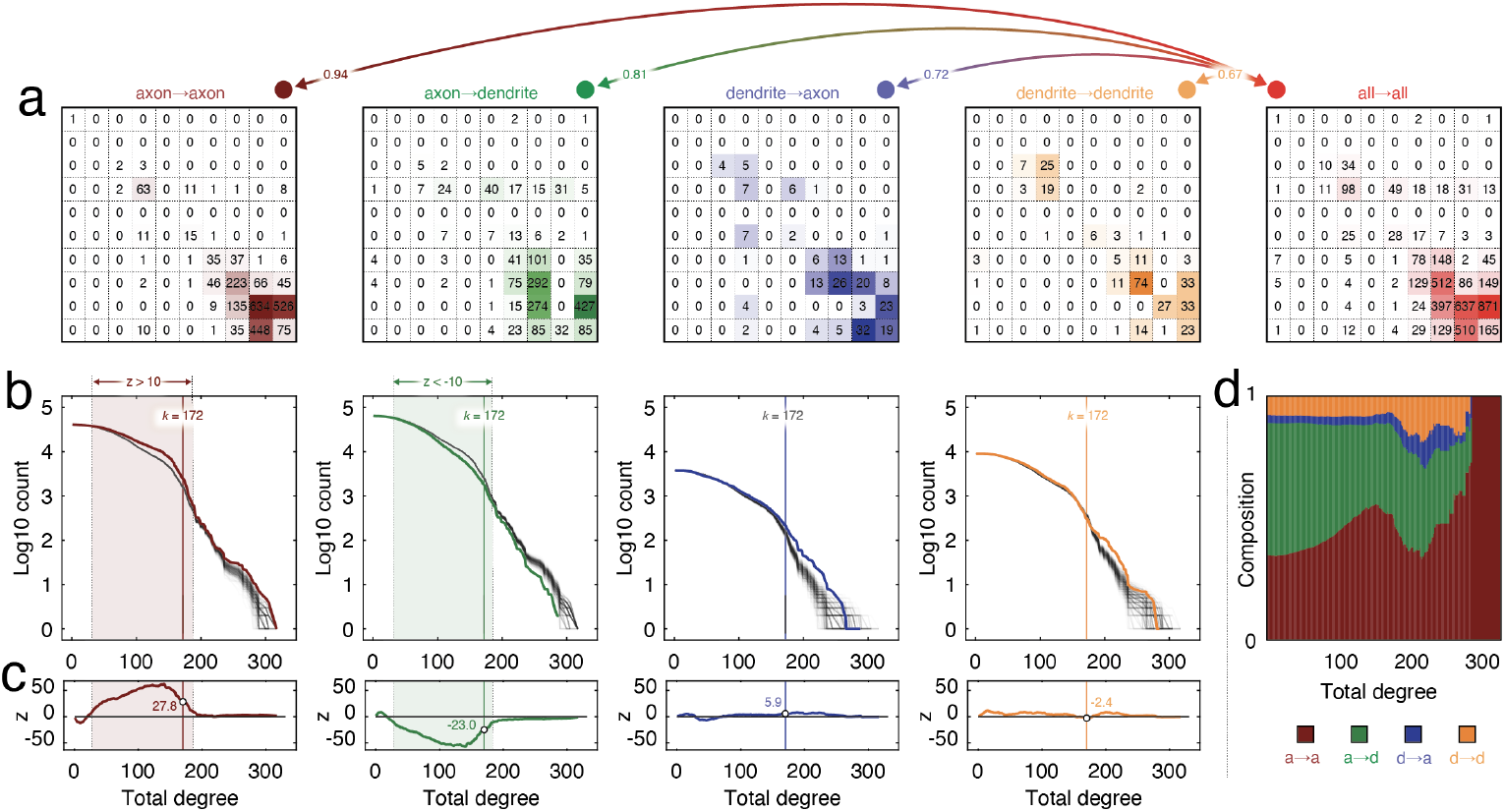
Linking rich-club organization to synapse type. In the main text we described the detection of a rich-club in the larval *Drosophila* connectome. This rich-club was defined using a composite connectivity matrix that combined four different synapse types: axon→axon, axon→dendrite, dendrite→axon, and dendrite→dendrite. Here, we break down the contributions of these different synapse types to the rich-club. Specifically, for the rich-club at *k* = 172, we calculate how “rich” connections–synapses that link rich-club members to other rich-club members–are distributed across the 10 level-4 communities. Panel *a* shows the binary connection counts between each of the 10 communities for each of the four synapse types. On the far right is the all→all matrix, which was reported in the main text and corresponds to the total connection count across all four synapse types. In general, we find that specific synapse types are similarly configured with respect to communities as that of the composite matrix. All four synapse types were significantly positively correlated with the all→all matrix (maximum correlation of *r* = 0.94; minimum correlation of *r* = 0.67), suggesting that different synapse types contribute to the rich-club with similar patterns. Next, we asked whether different synapse types were overrepresented in the rich club compared to their baseline frequency. To do this, we constructed the following null model. We constructed a “connectivity mask” that included all pairs of neurons connected by at least one of the four synapse types. Each connected pair of neurons was associated with a 4 *×* 1 vector whose elements denoted the number of axon→axon, axon→dendrite, dendrite→axon, and dendrite→dendrite synapses by which those neurons were connected. We then permuted the features vectors randomly across connected pairs of neurons. This null model preserves the baseline synapse counts across all four connection types, as well as the “connectivity mask”. That is, connected neurons in the empirical connectome remained connected in the null model. The only component that changed is the specific connection types by which those neurons were connected. We then generated 1000 surrogate networks using this null model and compared the observed connection type counts against those of the null model. We found that the axon→axon and axon→dendrite connections were over-/under-represented in the rich-club compared to their baseline prevalence. In more detail, the actual number of axon→axon and axon→dendrite rich-club connections were 2457 and 1769, respectively. Under the null model, we expected 1587*±*31 and 2486*±*31. This translates to z-scores of 27.81 and -23.0. Dendrite→axon and dendrite→dendrite connection were also over-/under-represented, but not nearly as extreme (z-scores of 5.9 and -2.4, respectively). Finally, we repeated these analyses at different degree cutoffs for defining the rich-club. We find that as *k* varies, the relative proportion of connection types changes. At low degree (where the rich-club is not well-defined) the composition reflects baseline connection-type prevalence. However, as degree increases and the rich-club becomes more exclusive (this range also includes the *k* = 172 rich-club reported in the main text), we find evidence that rich-club composition changes. Collectively, these results present a more nuanced view of the rich-club than what was reported in our initial submission. Specifically, we find that the rich-club is dominated by axon→axon and axon→dendrite connections. On one hand this is expected given that these connection types are most common. However, when we compare these counts against what would be expected under a null model, we find that the number of axon→axon rich-club connections is far greater than expected and, conversely, the number of axon→dendrite connections is far less than expected. These observations suggest that different connection types differentially contribute to the rich-club.

**Figure S13.**
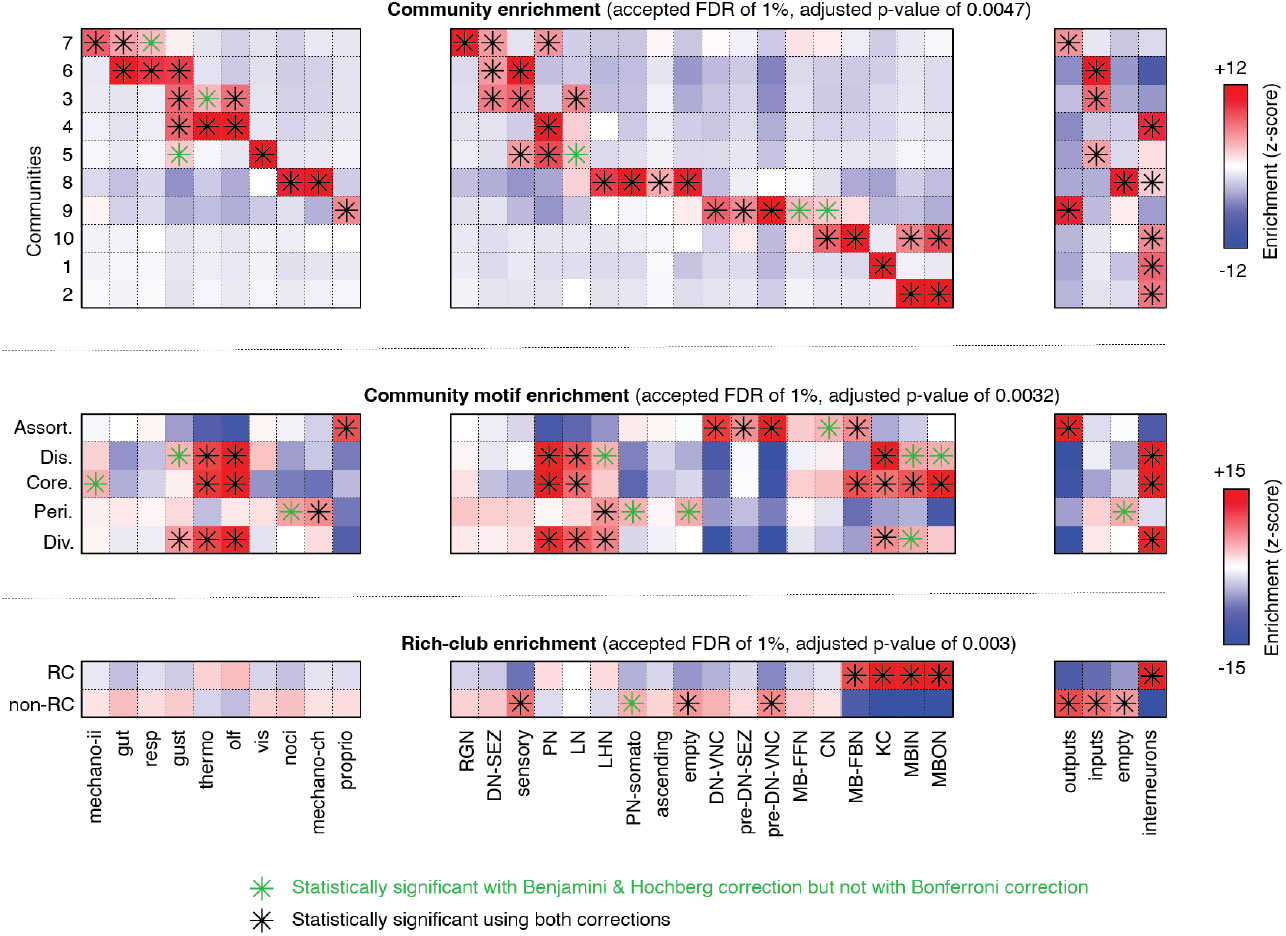
Comparison of strategies for controlling false discovery. In the main text we compared neuronal annotations with community labels, community motifs, and rich-club assignments. To correct for multiple comparisons, we followed the strategy of Benjamini and Hochberg [121], wherein we specify acceptable false discovery rate (1%, in every case here) and adjust the critical *p*-value so as to ensure that this tolerance is rate is not exceeded. However, given the nestedness of the hypotheses we test–e.g. the “functional” annotations are all subsets of the “sensory” cell type–it is unclear whether this procedure is effective. Here, we compare the results reported in the main text with the results obtained using a more conservative statistical criterion, namely the Bonferroni adjustment, wherein our original critical value is penalized by the number of tests. Here we set our critical value equal to *α* = 0.01 and obtain a Bonferroni-corrected critical value as 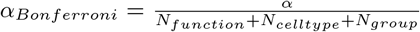, where *N*_*function*_, *N*_*celltype*_, and *N*_*group*_ are the number of tests. For example, when we compare communities to functional, cell type, and group annotations, the total number of tests is *N*_*total*_ = (10 *×* 10) + (10 *×* 18) + (10 *×* 4) = 320, resulting in a corrected critical value of 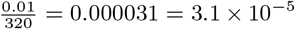. Here we show enrichment scores for community labels, community motifs, and rich-club assignment, highlighting values that are statistically significant under the Benjamini & Hochberg method (black and green asterisks) *versus* those that achieve statistical significance only under the Bonferroni correction (black asterisks). We note, however, that these two correction methods identify similar patterns of enrichment.

**Figure S14.**
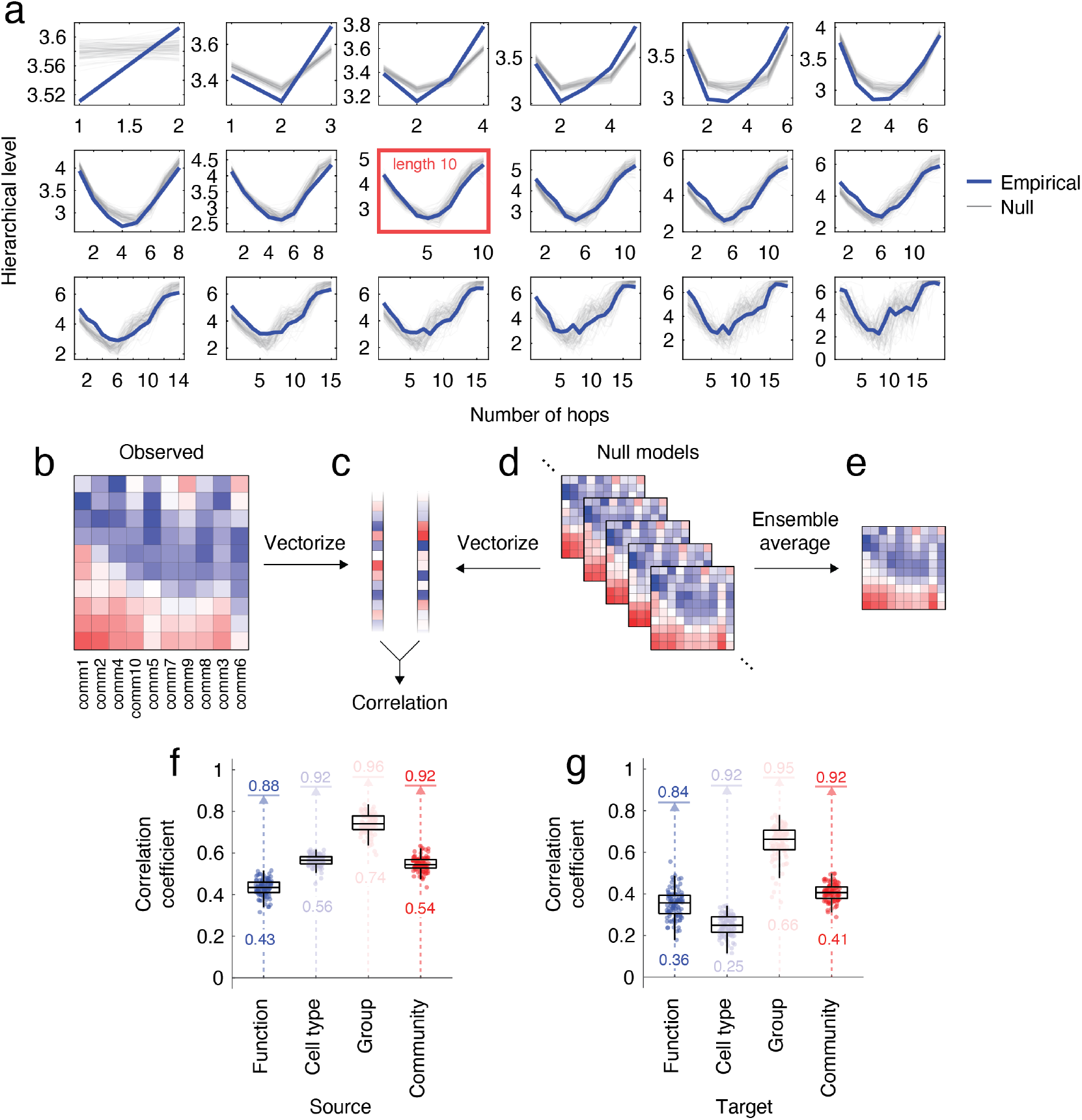
Comparing shortest paths trajectories against null model. In the main text we showed that shortest paths follow characteristic trajectories through the community hierarchy. The first and final steps use within-community edges and intermediate steps use progressively more between-community edges. Here, we compare the empirical results against those obtained from a null model. Specifically, we generated 100 instances of binary null networks from the fitted stochastic blockmodel. We supplemented these networks until their total number of edges matched that of the empirical network and their block-wise edge counts were also exact matches. We then added weights to edges in each block by randomly assigning the empirical weights to the edges in that block. Subsequently we calculated shortest paths between all pairs of nodes. (*a*) Although there were subtle differences, the null model exhibited the same “u-shaped” trajectory as the empirical data. This result holds across shortest paths of different lengths (hop counts). Each sub-panel within panel *a* shows trajectories for shortest paths of different lengths. In the main text we showed that these trajectories were specific to different functions, cell types, group types, and community labels. Here, we follow the same procedure as in the main text and derive typical trajectories for when each annotational label is either a source or target in a shortest path. We do so for the observed network and for each of the 100 null networks. After repeating the procedure for the null networks, we also calculate the ensemble average. We then vectorize the trajectories and compute their spatial similarity (correlation). Panels *b*-*e* illustrate this procedure using “community labels” as the example annotations. As in the main text, we focus on paths of length 10 (number of hops) but note that results generalize to other lengths. Panels *f* and *g* summarize the results. The boxplots show similarity of the empirical trajectories to each of the 100 null networks. The text below each boxplot is the median value. The arrows denote similarity to the ensemble average, which is also indicated by the accompanying text.

